# Emerging SARS-CoV-2 Variants of Concern: Spike Protein Mutational Analysis and Epitope for Broad Neutralization

**DOI:** 10.1101/2021.12.17.473178

**Authors:** Dhiraj Mannar, James W. Saville, Zehua Sun, Xing Zhu, Michelle M. Marti, Shanti S. Srivastava, Alison M. Berezuk, Steven Zhou, Katharine S. Tuttle, Michele D. Sobolewski, Andrew Kim, Benjamin R. Treat, Priscila Mayrelle Da Silva Castanha, Jana L. Jacobs, Simon M. Barratt-Boyes, John W. Mellors, Dimiter S. Dimitrov, Wei Li, Sriram Subramaniam

## Abstract

Mutations in the spike glycoproteins of SARS-CoV-2 variants of concern have independently been shown to enhance aspects of spike protein fitness. Here, we report the discovery of a novel antibody fragment (V_H_ ab6) that neutralizes all major variants, with a unique mode of binding revealed by cryo-EM studies. Further, we provide a comparative analysis of the mutational effects within variant spikes and identify the structural role of mutations within the NTD and RBD in evading antibody neutralization. Our analysis shows that the highly mutated Gamma N-terminal domain exhibits considerable structural rearrangements, partially explaining its decreased neutralization by convalescent sera. Our results provide mechanistic insights into the structural, functional, and antigenic consequences of SARS-CoV-2 spike mutations and highlight a spike protein vulnerability that may be exploited to achieve broad protection against circulating variants.

## Introduction

Genomic surveillance of SARS-CoV-2 during the first year of the COVID-19 pandemic revealed that the D614G mutation in the spike glycoprotein (S protein) was the sole widespread consensus mutation, with the G614 genotype largely replacing the D614 genotype in February 2020^1,2^. In November 2020 however, the emergence of the Alpha (B.1.1.7) variant began capturing global headlines and coincided with a surge in COVID-19 cases in the United Kingdom. Within 4 months, the Alpha variant became the globally dominant SARS-CoV-2 lineage^1^. The emergence of the Alpha lineage was quickly followed by the emergence of the Beta (B. 1.351), Gamma (P.1), and Epsilon (B. 1.427/429) variants in early 2021, with the Kappa and now globally dominant Delta variants emerging shortly thereafter.

Initial case tracking data demonstrated increased transmissibility for the Alpha, Beta, and Gamma variants of concern (VoC), with early biochemical data suggesting that increased ACE2 affinity imparted by the N501Y mutation to largely underlie the observed increase in infectivity^3^. Common amongst these variants are identical (N501Y, E484K) or similar (K417T, K417N) mutations within the receptor-binding domain (RBD), which may impact both ACE2 binding and antibody neutralization escape^4–6^. In addition, distinct mutations within highly antigenic loops in the N-terminal domain (NTD) across these variants primarily reduce antibody neutralization^7^.

Here, we present a novel antibody (ab6) with neutralization activity against multiple variants (Alpha, Beta, Gamma, Delta, Kappa, Epsilon) and report its epitope within the RBD using cryogenic electron microscopy (cryoEM). This antibody epitope is remote from most VoC mutations, explaining its ability to confer pan-variant neutralization. Given the enhanced antibody escape of circulating variant spikes, the epitope we define here provides opportunities for rational therapeutic targeting of variant SARS-CoV-2 S proteins. We also report studies of spike structure, ACE2 affinity, and evasion of antibodies afforded by variant spikes, providing a general structural rationale for enhanced viral fitness of the variants.

## Results

### Broad Neutralization of the SARS-CoV-2 Spike Protein by an Unconventional Antibody Fragment

V_H_ ab6 is a phage-display derived antibody with the unusual biochemical property of exhibiting enhanced RBD affinity as a monomeric fragment when compared to a bivalent fusion^8^, and was recently shown to exhibit tolerance to several circulating RBD mutations^9^. We first confirmed this anomalous property of ab6, showing that the bivalent V_H_-Fc fusion has lower neutralization potency relative to the monovalent V_H_ construct in both pseudotyped and live virus neutralization assays (Figure 1A). We then assessed V_H_ ab6 binding and neutralization of variant spikes, including the Delta and Kappa variant spikes (Figure 1B). Ab6 bound and neutralized all variant spike pseuodtyped viruses, but exhibited lower potency for the Epsilon, Kappa, and Delta spikes. We additionally assessed ab6 neutralization of Alpha, Beta, and Delta live viruses via authentic virus neutralization assays, confirming the lower potency of ab6 for the Delta variant. (Figure S1).

**Figure 1.**
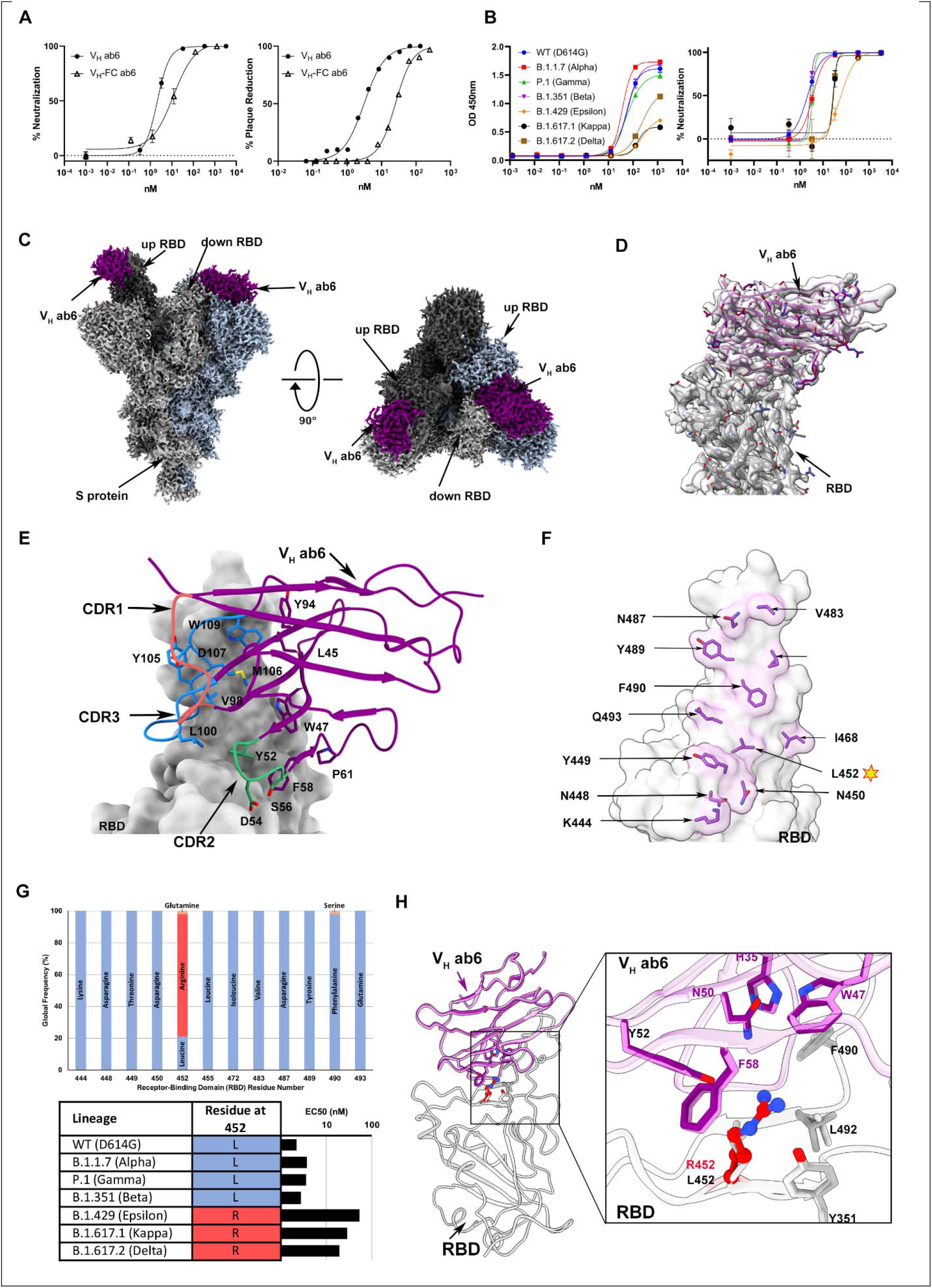
Ab6 broadly neutralizes SARS-CoV-2 variants via a largely conserved molecular epitope. **(A)** Pseudovirus neutralization (left) and authentic SARS-CoV-2 plaque reduction neutralization test - PRNT (right) by V_H_ ab6 and V_H_-Fc ab6 against wild-type lineages. The pseudovirus neutralization (left) utilized the D614G wild-type, whereas the PRNT (right) utilized the USA-WA1/2020 isolate. **(B)** ELISA S protein binding (left) and pseudovirus neutralization (right) of SARS-CoV-2 variants by V_H_ ab6. **(C)** 2.4Å global cryoEM density map of V_H_ ab6 bound to wild-type S protein. Density corresponding to S protein protomers and ab6 are shown in grayscale and navy blue, respectively. **(D)**Focus-refined atomic model of ab6 bound to the wild-type (D614G) S protein RBD. **(E)** Analysis of ab6 contact zones. The RBD and ab6 are shown as a gray molecular surface and colorized cartoon representation, respectively. The ab6 scaffold is depicted in purple and the three complementarity determining regions (CDRs) of ab6 are coloured as follows: CDR1 - red; CDR2 - green; CDR3 - blue. **(F)**Footprint of ab6 mapped onto the wild-type S protein RBD. The sidechains of footprint residues are shown and colorized in purple. **(G)** (Top) Global frequency of residue identity within the ab6 footprint in GISAID deposited sequences as of st August 1, 2021. (Bottom) Residue identity at position 452 within SARS-CoV-2 variants and associated V_H_ ab6 EC50 measured via pseudoviral neutralization assays depicted on a logarithmic scale. **(H)** Focused view superpositions of the cryoEM-derived atomic model of the Epsilon (B.1.429) and wild-type (D614G) S proteins bound to V_H_ ab6. Epsilon and wildtype RBD’s are coloured light and dark grey respectively, while purple and pink models refer to ab6-WT and ab6-Epsilon, respectively. The R452 mutation is highlighted in red. Pseudoviral neutralization experiments and ELISAs were performed in triplicate, error bars denote the standard error of the mean. PRNT experiments were performed in duplicate, and the mean is plotted.

We next determined the cryoEM structure of V_H_ ab6 bound to the WT spike at 2.57Å (Figure S2), showing that ab6 binds to the RBD in both the up and down positions, via a unique binding mode (Figure 1C). Local refinement enabled visualization of the ab6-RBD interface at 3.21Å (Figure S2G-H), and revealed that the ab6-RBD interaction is dominated by contacts with the ab6 beta-sheet scaffold, which wraps around the RBD, extending this large interface to include its CDR2 and CDR3 loops but leaving the CDR1 loop free (Figure 1E). This scaffold-mediated interaction necessitates a near perpendicular angle of approach for ab6 relative to the RBD (Figure 1E), which can only be accommodated by a single V_H_ within a bivalent fusion construct, and likely accounts for the unusual behavior of the V_H_-Fc ab6 construct relative to V_H_ ab6 (Figure 1A).

The ab6 footprint overlaps that of ACE2, consistent with its mechanism of neutralization via ACE2 competition^8^ (Figure S3). Analysis of the ab6 footprint reveals inclusion of L452, consistent with the reduced potencies against the Epsilon, Delta, and Kappa spikes, which harbour the L452R mutation (Figure 1F,G). Importantly, L452 is the only residue mutated in variant spikes that falls within the ab6 footprint, rationalizing its broad neutralization of all variants tested. Furthermore, analysis of genomic sequences from the GISAID database confirms the conserved nature of the ab6 epitope, highlighting the L452R mutation to be the only significantly occurring variation (Figure 1G). To uncover the structural basis for the attenuation of ab6 potency by the L452R mutation, we obtained the cryoEM structure of the Epsilon spike bound to ab6. A global 3D reconstruction was obtained at 2.45Å (Figure S4), in which ab6 bound both up and down RBDs, as seen in the WT-ab6 complex. Focused refinement enabled visualization of the ab6-Epsilon spike interface at 3Å (Figure S4G-H), revealing R452 to extend towards the ab6 scaffold (Figure 1H). This orientation places the positively charged R452 sidechain in close proximity to a hydrophobic patch within ab6, centered around F58. Thus, the reduced potency observed for R452 containing spikes is likely a result of unfavorable charge and steric effects.

To contextualize the broad-spectrum activity of ab6 we sought to investigate variant spike mutational effects on antibody evasion, receptor engagement, and spike structure. As we have previously reported these analyses on the Kappa and Delta variant spikes^10^, we proceeded here with our analysis of the Alpha, Beta, Gamma, and Epsilon variant spike proteins.

### RBD and NTD Directed Antibodies are Escaped by Variant Spike Proteins

Having demonstrated the broad neutralization of variant spikes by ab6, we next aimed to provide a comparative analysis of spike mutational effects and antibody breadth using a representative panel of previously reported monoclonal antibodies. We selected RBD directed antibodies^11–14^ which cover the four distinct anti-RBD antibody classes^15^ and an ultrapotent antibody, S2M11^16^, which uniquely binds two neighbouring RBDs simultaneously (Figure 2A). We additionally included the NTD-directed antibodies 4-8 and 4A8 to investigate the impact of NTD mutations within these variant spikes. (Figure 2A). Antibody binding was quantified via enzyme-linked immunosorbent assay (ELISA), and compared with neutralization, which was measured via a pseudoviral entry assay (Figure 2B). S309 and CR3022 are cross-reactive SARS-CoV-1 directed antibodies whose footprints do not span VoC mutations, and accordingly exhibited relatively unchanged binding across all variant spikes. We have previously characterized the mutational sensitivity of ab1, ab8, and S2M11 to spikes bearing only RBD mutations, and the current analysis of antibody evasion using spikes bearing all VoC mutations is consistent with our previous report^4^: 1) The N501Y mutation within the Alpha variant reduces but does not abolish the potency of ab1, while dramatic loss of ab1 activity is seen in Beta and Gamma variants due to mutation of K417 to N or T, respectively; 2) The E484K mutation abrogates ab8 activity in the Beta and Gamma variants; and 3) the L452R mutation reduces but does not abrogate activity of S2M11 in the Epsilon variant spike, drawing similarity to the mutational sensitivity of ab6. To compare the structural basis for the effect of L452R on S2M11 and ab6, we performed cryoEM studies on the Epsilon-S2M11 complex (Figure S6). We obtained a global 3D reconstruction of the Epsilon spike bound to three S2M11 Fabs at 2.16Å. In contrast to the Epsilon-ab6 structure wherein which R452 protrudes into the antibody interface (Figure 1H), R452 extends away from the S2M11 interface (Figure S7). As the L452 sidechain is accommodated within the footprint of S2M11, this positioning of R452 is likely due to steric clashing and charge repulsion effects which may increase the interaction energy and underlie the observed attenuation of potency.

**Figure 2.**
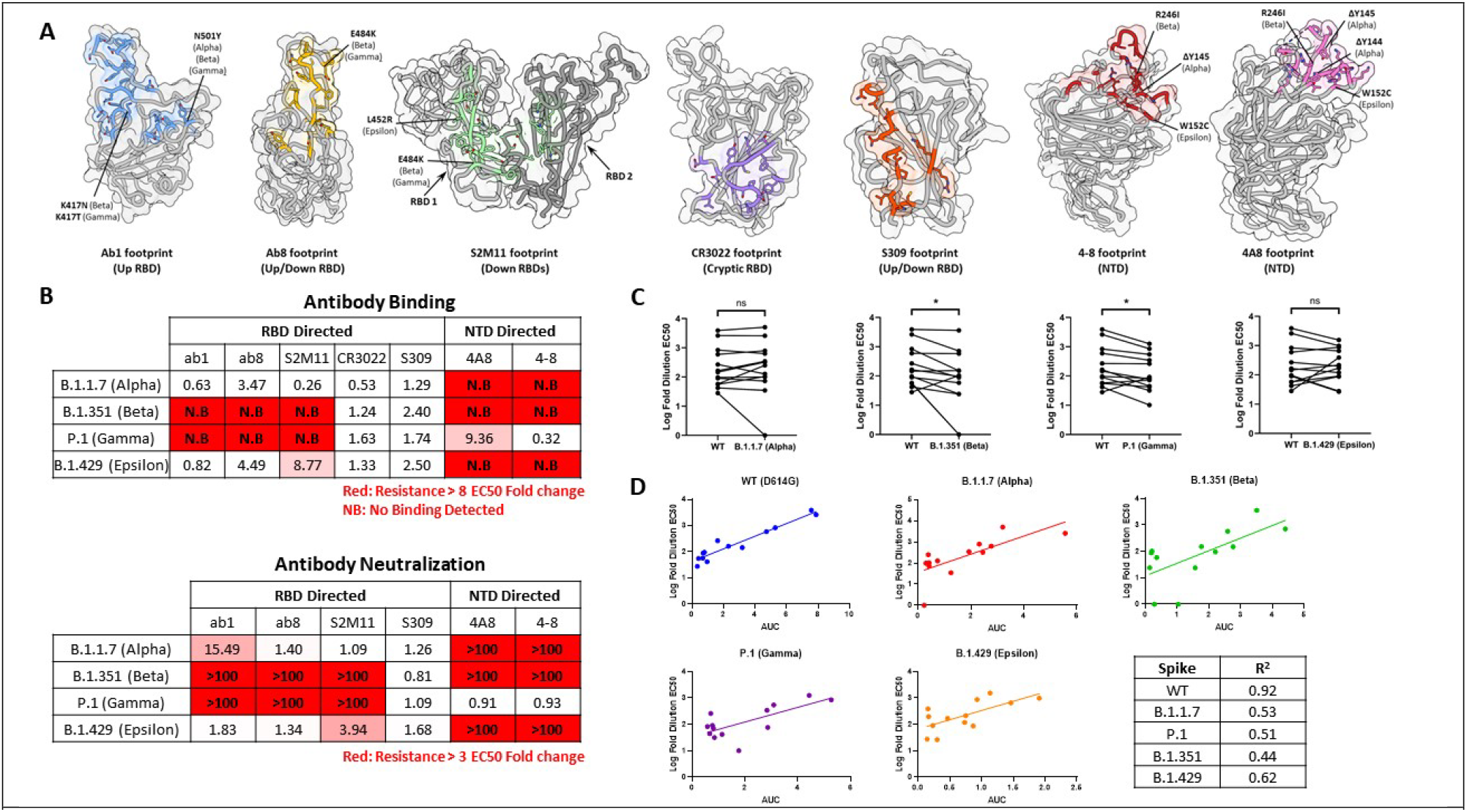
Alpha, Beta, Gamma, and Epsilon S proteins exhibit differences in monoclonal and polyclonal antibody escape. **(A)** Antibody binding footprints for monoclonal antibodies included in the study. Variant S protein mutations falling within each footprint are highlighted. **(B)** Fold-changes in antibody binding (top) and pseudovirus neutralization EC50s (bottom) for each variant spike relative to wild-type (D614G). Antibody binding was quantified by ELISA. **(C)** Log-fold EC50 dilutions of patient sera when neutralizing wild-type and variant spike pseudovirus. Statistical significance was tested via the Wilcoxon matched pairs test (*P ≤ 0.05, ns, not significant). See figure S8 for patient information. **(D)** Correlation between NTD+RBD binding antibody levels and neutralization of wild-type and variant pseudoviruses. (AUC = area under the curve of the NTD+RBD ELISA binding curves). Correlation coefficients (R) are tabulated in the bottom right of panel (D).

Evasion of NTD-directed antibodies was observed in cases when mutations were either within, or adjacent to, antibody footprints (Figure 2B), corroborating the recently described remodelling of antigenic loops in the Alpha, Beta, and Epsilon spikes^17,18^. The W152C substitution within the Epsilon NTD is inside the footprints of 4A8 and 4-8, and both antibodies were escaped by this variant spike. The Beta NTD contains a deletion (Δ242-245) which spans the 4-8 footprint, along with the R261I substitution spanning both 4A8 and 4-8 footprints, leading to escape from both antibodies. The footprint of 4A8 and 4-8 spans a deleted site within the Alpha NTD (Δ144-145) leading to escape. These direct and allosteric mutational effects are consistent with previous findings on NTD rearrangement within these variants and demonstrate their antibody evasive properties.

Having characterized monoclonal antibody evasion by variant spikes, we extended our analysis to include polyclonal antibody escape from human sera. Sera were collected from a spectrum of patients with varying COVID-19 infection histories and vaccination statuses (Figure S8C) and subjected to neutralization and binding assays (Figures S8A-B, S9). Potent neutralization of WT spike pseudovirus was observed in all COVID-19 positive or vaccinated samples but not with pre-pandemic sera from uninfected patients, suggesting limited pre-existing immunity (Figure S8A-B). While serum levels of spike ectodomain binding antibodies correlated poorly with wildtype spike neutralization, strong correlations were observed between NTD and RBD binding antibody levels and neutralization (Fig S10), corroborating the dominance of neutralizing epitopes within the NTD and RBD^15,19,20^. We observed various effects on neutralization escape when sera samples were assayed using variant spike pseudotyped viruses, obtaining statistically significant decreases in neutralization efficacy for both Beta and Gamma variants relative to wild-type (Figure 2C). Interestingly, the high correlation between serum NTD and RBD binding antibodies and pseudovirus neutralization for wild-type spikes was markedly reduced for all variant spikes (Figure 2D). Taken together, these results highlight the role of mutations within the NTD and RBD of variant spikes in driving evasion of SARS-CoV-2 directed monoclonal and polyclonal antibodies, providing a basis to evaluate the broad mutational tolerance exhibited by S309 and ab6.

### Enhanced Receptor Binding by Variant Spike Proteins

In addition to driving antibody escape, variant spike mutations can enhance receptor engagement, which may underlie increases in infectivity. To investigate the ACE2 binding potential of SARS-CoV-2 variant spikes, recombinant S protein ectodomains bearing variant spike mutations were used in biolayer interferometry (BLI) experiments. All mutant spikes exhibited slightly higher affinities for immobilized dimeric Fc-ACE2 when compared to wildtype (Figure S11A-B). Additionally, we used flow cytometry to evaluate the ability for recombinant dimeric Fc-ACE2 to bind wild-type or variant full-length spikes which we transiently expressed in Expi-293 cells. We did not observe major differences in spike protein expression across the variants (Figure S12A), and all mutant spikes tested demonstrated marginally enhanced ACE-2 binding potencies relative to wild-type (Figure S11C). These complementary assays demonstrate that the totality of mutations within each variant S protein enable slightly enhanced ACE2 binding, suggesting a contributing factor for the increased infectivity observed for these SARS-CoV-2 variants.

### Structural Effects of Variant Spike Protein Mutations

Having demonstrated mutational effects on antibody evasion and receptor engagement, we next sought to characterize the structural impacts of variant S protein mutations. To this aim, ectodomains bearing variant spike mutations were used for cryoEM structural studies. Global 3D reconstructions were obtained at resolutions ranging from (2.25-2.5 Å) (Figure S13-16), yielding open trimers with one RBD in the up conformation and 2 RBD’s in the down conformation for all spikes (Figure 3). The resolution within the NTD and RBD was insufficient for accurate visualization of mutational impacts within these domains, due to high degrees of conformational heterogeneity. In contrast, we were able to confidently model sidechains close to, and within the S2 domain, owing to its limited flexibility. We therefore first focused our analysis on mutational effects within this region, which predominantly localized to inter-protomer interfaces.

**Figure 3.**
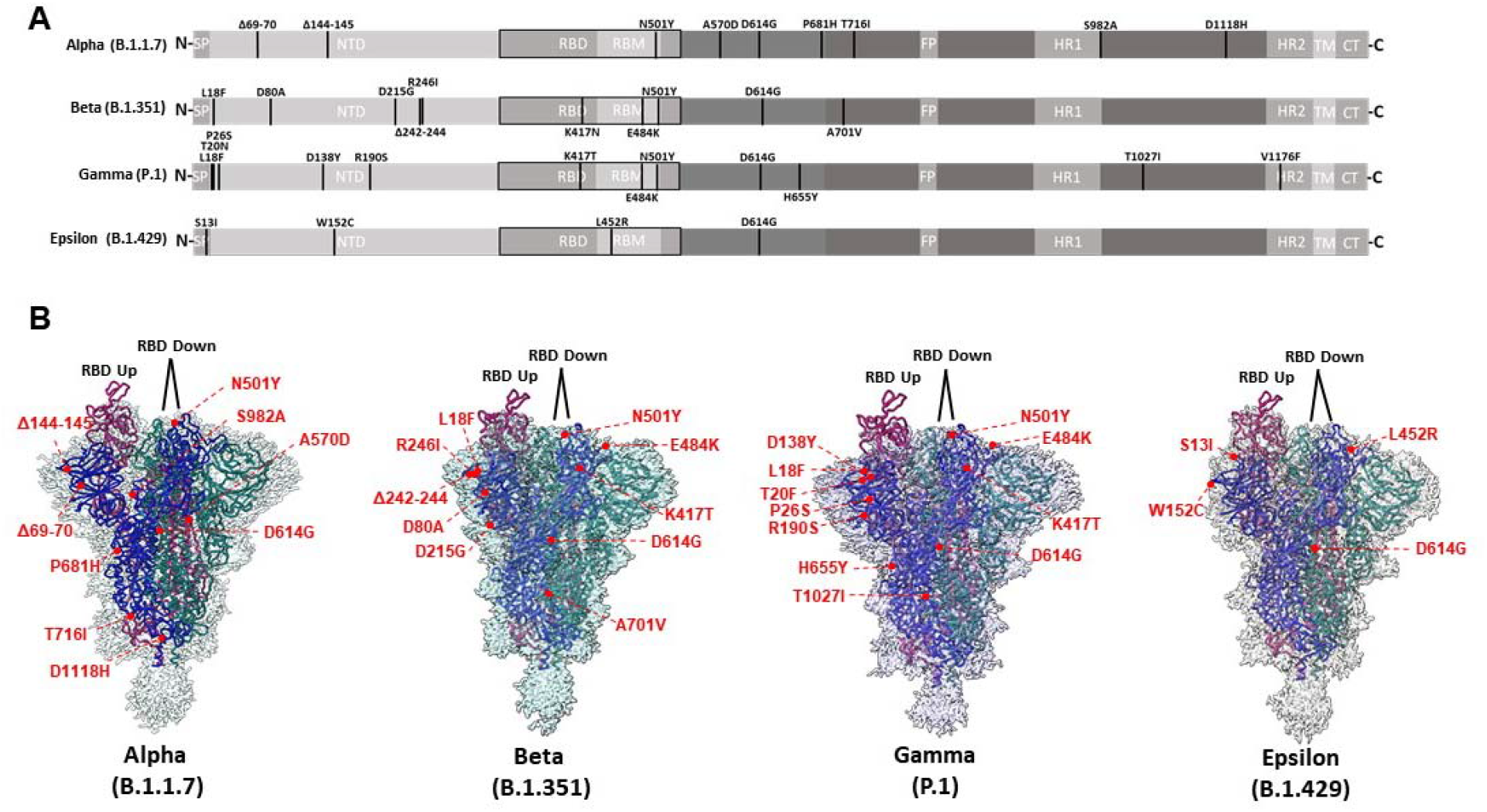
CryoEM structures of Alpha, Beta, Gamma, and Epsilon spike glycoproteins. **(A)** Linear schematic depicting mutations within variant S proteins (SP: Signal peptide, NTD: N-terminal domain, RBD: Receptor binding domain, RBM: Receptor binding motif, FP: Fusion peptide, HR1: Heptad repeat 1, HR2: Heptad repeat 2, TM: Transmembrane, CT: Cytoplasmic tail). **(B)** Global cryoEM maps and models for the Alpha (2.56 Å), Beta (2.56 Å), Gamma (2.25 Å), and Epsilon (2.4 Å) variant S proteins. Mutational positions are indicated and labelled in red.

Inspection of the structure of the Alpha variant shows that the A570D and S982A mutations appear to contribute protomer-specific structural effects with implications for RBD positioning (Figure 4B). Within both “RBD down” protomers, D570 either occupies a position within hydrogen bonding distance of N960 within the adjacent “RBD down” protomer, or sits within intra-protomer hydrogen bonding distance with T572 when the RBD of the adjacent protomer is in the up conformation (“RBD up”). D570 within the “RBD up” protomer uniquely forms a salt bridge with K854 in the adjacent “down RBD” protomer. S982 sits within hydrogen bonding distance of residues G545 and T547 within adjacent “RBD down” protomers only, and such interactions are not possible with the S982A mutation. Thus, the A570D and S982A mutations likely modulate RBD conformation through 1) stabilizing “RBD up” protomers via addition of the K852-D570 salt bridge, and 2) disruption of “RBD down”protomers via loss of stabilizing interactions afforded by S982.

**Figure 4.**
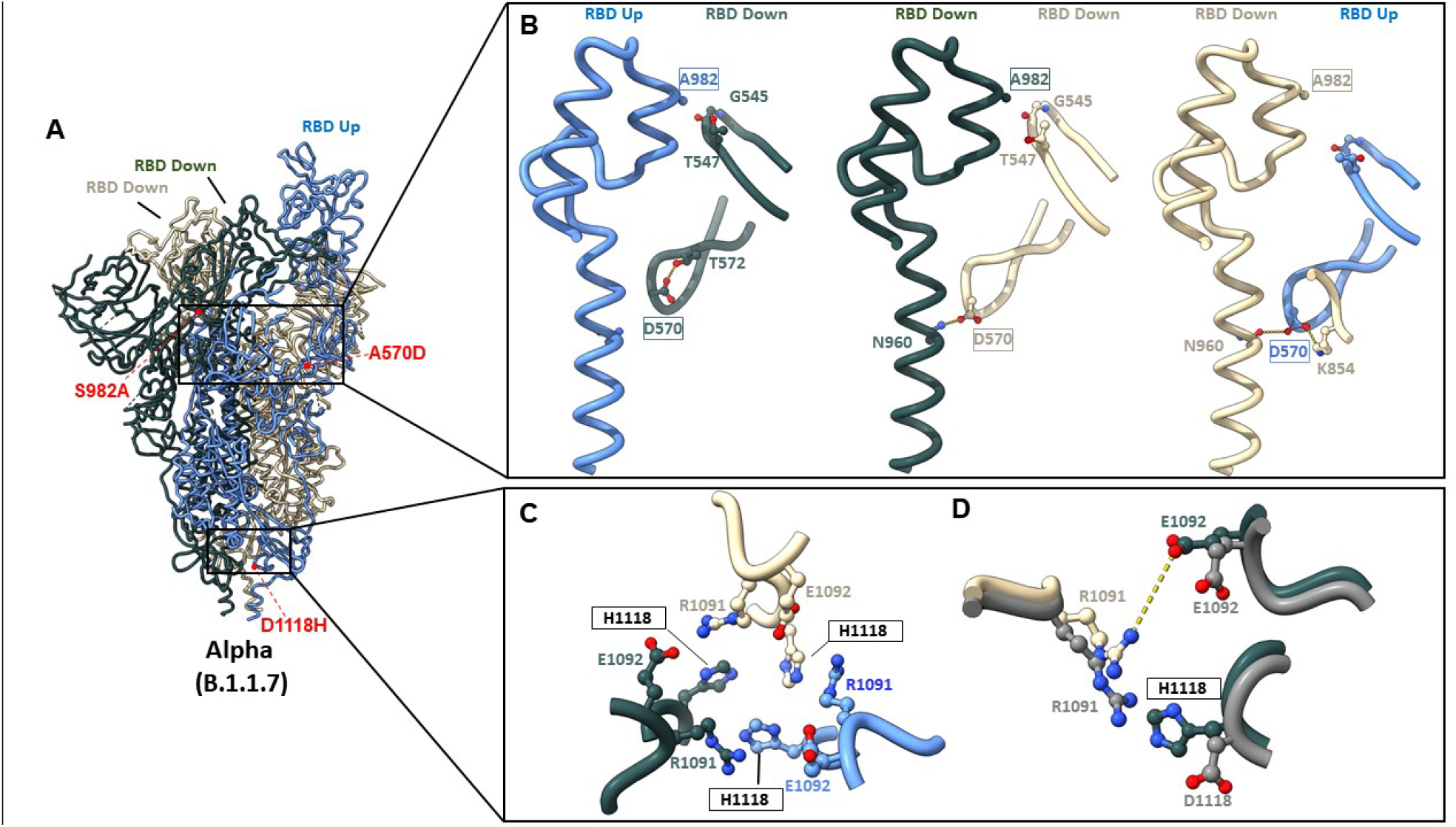
Alpha variant S protein mutations result in structural changes at inter-protomer interfaces. **(A)** Global model of the Alpha variant S protein with one RBD in the “up” conformation (blue) and two RBDs in the “down” conformation (beige and grey). Relevant mutations at inter-protomer interfaces are highlighted in red. **(B)** Modulation of inter-protomer contacts by mutations S982->A and A570->D between protomers with differential RBD conformations. **(C,D)** Impact of the D1118->H mutation on inter-protomer contacts. (C): model of H1118 and adjacent residues within the Alpha spike. (D): Superposition demonstrating differences between the Alpha spike (colored) and the wild-type spike (PDB: 6VSB) (shown in grey). Potential interactions (either electrostatic or hydrogen bonding) are highlighted as dashed lines. Mutated residues are indicated as boxed labels.

The D1118H mutation within the Alpha variant enables local side chain rearrangements, giving rise to additional interprotomer contacts, via pi-cation interactions between R1091 and H1118 of adjacent protomers, and electrostatic interactions between R10192 and adjacent E1092 residues (Figure 4C). Superposition of wild-type and Alpha spike models clearly demonstrates the differential positioning of H1118 compared to D1118, and the resulting movement of adjacent R1091 towards E1092 (Figure 4D). These additional interprotomer contacts enabled by the D1118H mutation may aid in the stabilization of the Alpha spike protein in its trimeric form.

Additional mutations were visualized within the Beta and Gamma variant spike proteins, with implications on inter-protomer and intra-protomer contacts. The A701V mutation in the Beta variant S protein lies at the protein’s surface and the larger sidechain conferred by V701 may enable tighter interprotomer packing with Q787 on adjacent protomers (Figure S17A). The T1027I mutation lies within the Gamma spike S2 core and the bulky hydrophobic I1027 sidechain faces inwards, increasing the hydrophobicity of the local buried environment (Figure S17B). This may decrease the local dielectric constant, enhancing the strength of the nearby network of interprotomer salt bridges between E1031 and R1039 on adjacent protomers (Figure S17B) and thus stabilize the Gamma variant trimer.

To identify structural mutational impacts on ACE2 binding we next determined the cryoEM structures of variant spike-ACE2 complexes. Resulting maps were obtained at average resolutions of ~2.6-3Å (Figures S18-S21). Focus-refined structures of the ACE2-RBD interface enabled visualization of RBD mutations, revealing local structures which are identical to our previously reported structures using spikes harbouring variant RBD mutations alone^4^. Superposition of local RBD-ACE2 complex models revealed no significant structural changes (Figure S22). These structural findings confirm that mutations outside of the RBD do not modulate positioning of ACE2-contacting residues at the receptor interface via allosteric mechanisms.

### Structure of the Gamma Variant NTD

Structure of NTD region in the Alpha, Beta, and Epsilon variants has been previously reported^17,18^. Here, we report structural analysis of the NTD region in the Gamma variant, stabilized using Fab fragments of the NTD-directed antibodies 4A8^21^ and 4-8^22^. The bound antibody fragments improve resolution of this flexible domain, enabling determination of the structure at resolutions of ~ 2.6 Å both for the overall spike and the NTD-antibody interface (Figures S23-S24).

Superposition of 4A8-bound wild-type and Gamma NTDs reveals remodelling of the antigenic supersite N1 loop^23^ within the Gamma variant but not the N3 and N5 loops, which comprise the majority of the 4A8 binding site (Figure 5B). Analysis of nearby mutational effects provides additional reasoning for conformational remodelling of the N1 loop. The mutations L18F, D138Y, and T20N cluster close together, forming multiple interactions which stabilize the alternate N1 conformation (Figure 5C). Namely, F18 and Y138 form an interconnected network of T-shaped pi stacking interactions with each other and the adjacent F140 residue. Additionally, Y138 and N20 sit within hydrogen bonding distance of the main chain carbonyl of F79 and the sidechain of D80, respectively. Comparison of sidechain positioning between wild-type and gamma structures in this region reveals steric clashes between Gamma residue Y138 and wild-type residues F79 and L18, and between Gamma residue N20 and the main chain of wild-type residue L18, resulting in differential positioning of D80 and F79 in the Gamma NTD (Figure 5D). Identical positioning of the N1 loop is observed in the Gamma NTD-4-8 structure, further confirming these mutational effects (Figure S25). Thus, the unique interactions conferred by mutations within the Gamma NTD stabilize local conformations which are sterically incompatible with wild-type N1 positioning, causing N1 loop rearrangement.

**Figure 5.**
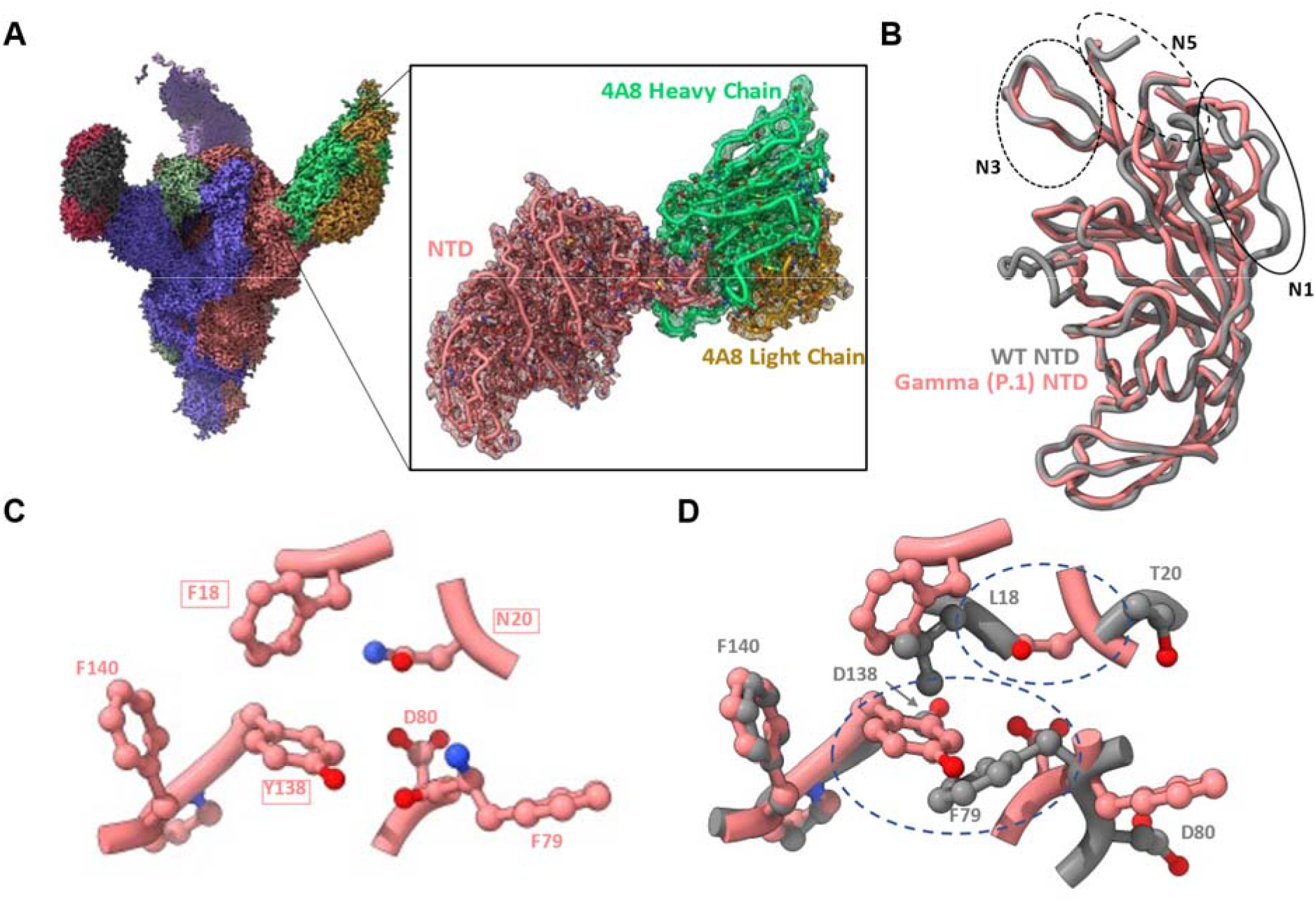
Structure of the Gamma variant NTD reveals rearrangement of the N1 loop. **(A)** Global cryoEM density map of the Gamma variant S protein bound to 4A8 at 2.59 Å (left), and focus-refined map and model for the Gamma NTD-4A8 complex at 2.66 Å (right). **(B)** Superposition of 4A8-bound Gamma and wild-type NTD models showing N1 loop rearrangement. The three loops (N1, N3, N5) comprising the “NTD neutralization supersite” are indicated with circles. **(C)** Positioning of the L18->F, D138->Y, and T20->N mutations and adjacent residues in the Gamma NTD. **(D)** Superposition of residues shown in (C) with WT residues demonstrates steric incompatibilities. Areas of steric clashes are indicated by dashe dovals. Mutated residues are indicated as boxed labels. The wild-type - 4A8 model (PDB: 7C2L) was used for all superpositions and is shown in grey throughout the figure.

## Discussion

Mutational enhancement of SARS-CoV-2 viral fitness can arise from effects on receptor engagement and evasion of neutralizing antibodies, with structural origins in the spike glycoprotein. Here we have examined these effects, demonstrating domain-specific differences in the roles and structural mechanisms of S protein mutations. Although such mutational changes can pose threats to natural and vaccine induced immunity, the existence of preserved epitopes within functional domains holds great potential for future antigenic focus. This is highlighted in our analysis of variant SARS-CoV-2 spikes, which despite exhibiting effects on antibody evasion and ACE2 binding, shared a conserved epitope within the RBD which conferred broad neutralization.

The structural impacts of VoC S protein mutations offer insight regarding the differing mutational heterogeneity observed for the NTD and RBD. While VoC mutations within the RBD are limited to only substitutions at a few residues (K417N/T, L452R, T478K, E484K/Q, N501Y), the NTD hosts a large array of deletions and substitutions, predominantly localizing to the three loops constituting the “NTD neutralization supersite” (N1: residues 14-26, N3: residues 141-156, N5: residues 246-260)^23^. Our structures of VoC S proteins in complex with ACE2 demonstrate minimal structural changes in the RBD, reflecting its functional constraints in cell-attachment, only permitting mutations that preserve the ACE2 binding interface. In contrast, our structure of the Gamma NTD confirms the role of mutations within this domain as enabling structural rearrangement of antigenic loops, a feature common to all variant spike NTDs (Alpha, Beta, Delta, Epsilon)^17,18^. These rearrangements are likely directed primarily by immune evasive pressures. Taken together, these contrasting structural effects between variant NTD and RBD mutations likely arise due to different functional requirements and selective pressures between these domains.

Analysis of mutational effects within the NTD, RBD and S2 domain enables a mechanistic understanding of domain-specific ramifications on individual aspects of S protein functionality. ACE2 and RBD-directed antibody binding analyses by variant spikes corroborated the simultaneous increase in receptor engagement and antibody evasion^4^, consistent with the functional role of the RBD. Significant NTD structural rearrangements as reported here and previously by others, in combination with antibody binding and neutralization assays, highlight antibody escape to be the major driver of NTD mutation. Our structural analysis of mutations within the S2 domain reveals effects on interprotomer contacts, suggesting evolutionary pressures on quaternary structure and stability. We found the A570D and S982A mutations in the Alpha spike protein to work in concert to modify promoter-promoter interactions allosterically modulate RBD positioning, a finding consistent with recent reports^17,24,25^. Additionally, the D1118H and T1027I mutations within the S2 core of the Alpha and Gamma spikes respectively, cause local changes which likely serve to enhance the strength of nearby inter-protomer salt bridges, with implications for spike stability. Therefore, the distinct mutational impacts observed for mutations occurring in the RBD, NTD, or S2 highlight the defined roles that these domains play in overall S protein function and likely reflect domain-specific mutational pressures.

Despite these mutational effects, several lines of evidence have emerged from the present study demonstrating the existence of pan-variant epitopes. The high correlation between RBD + NTD binding antibody levels and viral neutralization potency reflects the dominance of neutralizing epitopes within these domains of the WT spike (Figure 2D). The diminished correlation between these parameters when assessing variant spikes demonstrates mutational escape within these domains. However, the fact that neutralization of variant spikes is attenuated, but not abolished, suggests the preservation of neutralizing epitopes. The existence of such epitopes within the RBD is corroborated by the unaltered potency of the SARS-CoV-1 directed antibody S309 across Alpha, Beta, Gamma, and Epsilon spikes (Figure 2B). Most importantly, we reveal a novel epitope within the RBD conferring broad neutralization of all variant of concern spikes, including the rapidly spreading Delta variant, as evidenced by the broad neutralization spectrum of ab6. This epitope has largely survived viral evolution thus far, with the only prominent mutational change being L452R. This mutation is accommodated by ab6, albeit yielding decreased neutralization potencies. Therefore, we highlight this epitope for focus in the design of broadly protecting therapeutic antibodies and immunogens.

Several RBD mutation resistant antibodies against SARS-CoV-2 have been reported during the preparation of this manuscript^26–30^, providing additional context regarding the conserved epitope we report here. Antibodies DH1047^31^ and STE90-C11^28^ were isolated from convalescent patients, and SARS2-38^26^ from immunized mice. All three antibodies are RBD directed and bind epitopes distal to that of ab6 (Figure 6). While STE90-C11 tolerated most circulating RBD mutations, it exhibited loss of activity against the K417T, K417N, and N501Y mutations^28^, which are present in many VOC/VOI spike proteins. In contrast, SARS2-38 and DH1047 bind highly conserved epitopes, retaining potency across all VOC/VOI spikes, with DH1407 exhibiting cross reactivity with additional sarbecoviruses^26,30^. V_H_ ab6 is distinguished from these previously reported antibodies by its unique angle of approach and binding mode involving multiple V_H_ scaffold - RBD contacts (Figure 1E), along with its small (15 kDa) size. Small antibody fragments are attractive therapeutic modalities given their enhanced tissue penetration compared to conventional monoclonal antibodies^32,33^.

**Figure 6.**
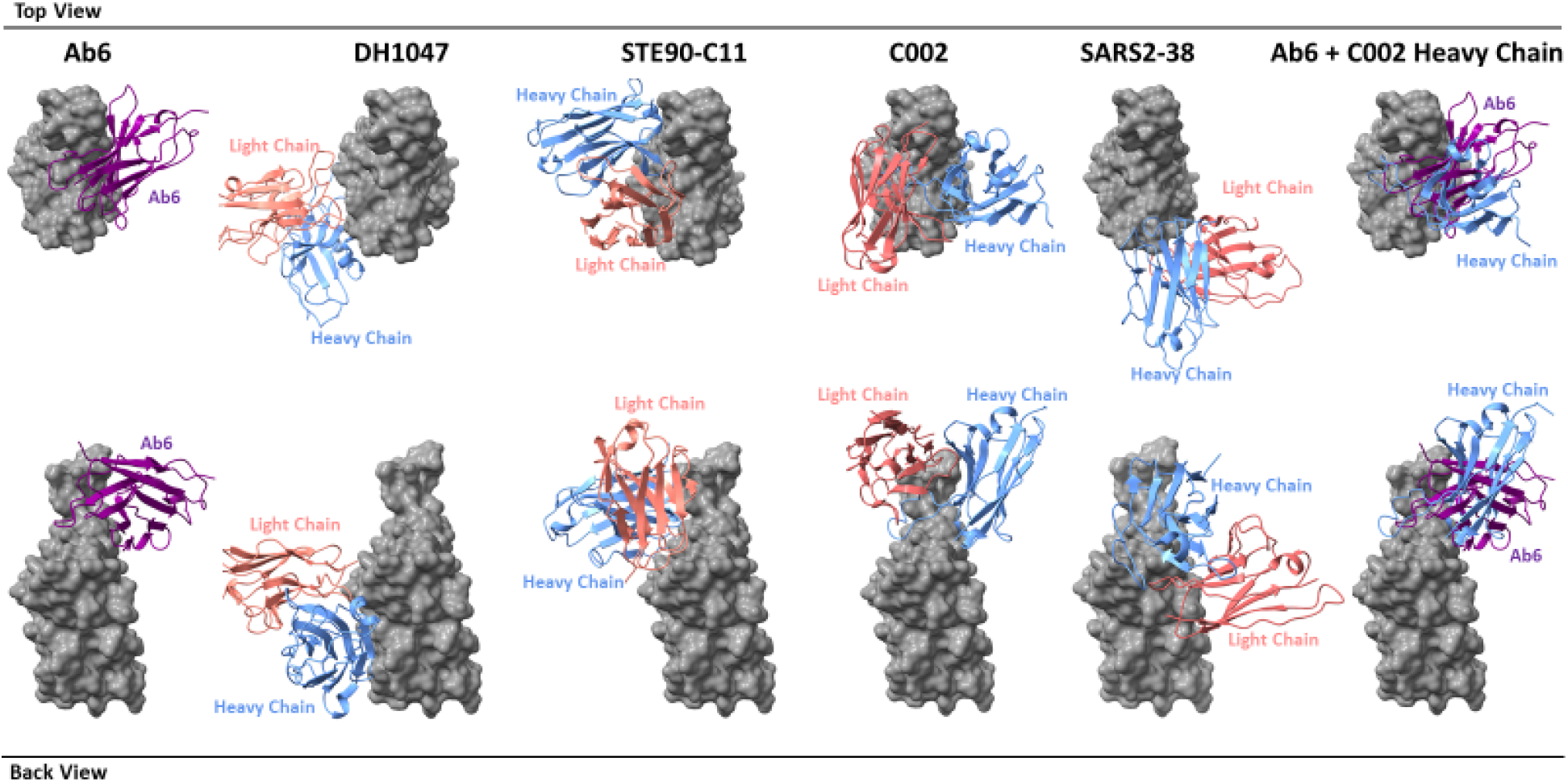
Footprint comparison between ab6 and selected RBD-directed antibodies. The RBD is depicted as a grey molecular surface and antibodies are depicted as colourized cartoon models. The following PDB files were utilized: 7LD1 (DH1047), 7B3O (STE90-C11), 7K8T (C002), 7MKM (SARS2-38). The RBD model from the ab6-RBD complex is shown for all antibody complexes for ease of visualization. For superpositions, structures were aligned using the RBD.

A recent study by a global consortium defined seven RBD binding antibody communities and showed broadly neutralizing antibodies either bind cryptic epitopes within the inner RBD face (communities RBD-6, RBD-7), or are non-ACE2 competing antibodies which bind the outer RBD face (community RBD-5)^29^. Ab6 binds the inner RBD face and contacts the RBM, enabling ACE2 competition, drawing similarity to the RBD-4 antibody community, which interestingly was not shown to contain any broadly neutralizing antibodies. Structural comparison of the ab6 footprint with a representative RBD-4 antibody (C002)^15^ reveals an overlapping footprint shared by the C002 heavy chain and ab6 despite differences in binding modes (Figure 6). C002 is derived from a convalescent patient, suggesting the potential for such an epitope to be recognized by natural antibodies. This evidence further supports the potential value of focus on the ab6 binding epitope for future therapeutic design.

## Materials and Methods

### Plasmids and Cloning

SARS-CoV-2 S protein Alpha, Beta, Gamma, and Epsilon genes were synthesized and inserted into pcDNA3.1 (GeneArt Gene Synthesis, Thermo Fisher Scientific). Alpha, Beta, Gamma, Epsilon, Delta, and Kappa mutated S protein ectodomain genes were synthesized incorporating the hexa-proline stabilizing mutations, the transmembrane domain replaced with a fibrin trimerization motif, and a C-terminal hexa-histidine tag (GeneArt Gene Synthesis, Thermo Fisher Scientific). The HexaPro expression plasmid was a gift from Jason McLellan (Addgene plasmid #154754; http://n2t.net/addgene:154754; RRID:Addgene_154754) which we used to construct the D614G mutant via site directed mutagenesis (Q5 Site-Directed Mutagenesis Kit, New England Biolabs).

C-terminal 7x his tagged NTD (amino acids 1-305) and RBD (amino acids 319 – 541) constructs were PCR amplified from D614G, Alpha, Beta, Gamma, and Epsilon full length spike ORFs. NTD constructs were cloned into pcDNA 3.1 using Nhel and Mssl restriction enzyme cloning, while the RBD constructs were introduced in frame to the mu phosphatase signal sequence and incorporated within pcDNA3.1 via Gibson assembly (NEBuilder HiFi DNA Assembly Cloning Kit, New England Biolabs).

Human ACE2 (residues 1-615) with a C terminal 7x his tag was amplified from “hACE2”, a kind gift from Hyeryun Choe (Addgene plasmid # 1786) and cloned into pcDNA3.1 via BstXI and XbaI restriction enzyme cloning. Successful cloning was confirmed by Sanger sequencing (Genewiz, Inc.) for all constructs.

### Pseudovirus Neutralization Assay

Variant and D614G spike pseudotyped retroviral particles were produced in HEK293T cells as described previously^34^. Briefly, a third-generation lentiviral packaging system was utilized in combination with plasmids encoding the full-length SARS-CoV-2 spike, along with a transfer plasmid encoding luciferase and GFP as a dual reporter gene. Pseudoviruses were harvested 60 h after transfection, filtered with a 0.45 μm PES filter, and frozen. For neutralization assays, HEK293T-ACE2-TMPRSS2 cells^35^ (BEI Resources cat# NR-55293) were seeded in 96-well plates at 50 000 cells. The next day, pseudovirus preparations normalized for viral capsid p24 levels (Lenti-X^™^ GoStix^™^ Plus) were incubated with dilutions of the indicated antibodies or sera, or media alone for 1 h at 37°C prior to addition to cells and incubation for 48 h. Cells were then lysed and luciferase activity assessed using the ONE-Glo^™^ EX Luciferase Assay System (Promega) according to the manufacturer’s specifications. Detection of relative luciferase units was carried out using a Varioskan Lux plate reader (Thermo Fisher). Percent neutralization was calculated relative to signals obtained in the presence of virus alone for each experiment.

### Expression and Purification of Recombinant Spike Proteins

Expi293F cells (Thermo Fisher, Cat# A14527) were grown in suspension culture using Expi293 Expression Medium (Thermo Fisher, Cat# A1435102) at 37°C, 8% CO_2_. Cells were transiently transfected at a density of 3 x 10^6 cells/mL using linear polyethylenimine (Polysciences Cat# 23966-1). The media was supplemented 24 hours after transfection with 2.2 mM valproic acid, and expression was carried out for 3–5 days at 37°C, 8% CO_2_. For ectodomain trimer purification, the supernatant was harvested by centrifugation and filtered through a 0.22-μM filter prior to loading onto a 5 mL HisTrap excel column (Cytiva). The column was washed for 20 CVs with wash buffer (20 mM Tris pH 8.0, 500 mM NaCl), 5 CVs of wash buffer supplemented with 20 mM imidazole, and the protein eluted with elution buffer (20 mM Tris pH 8.0, 500 mM NaCl, 500 mM imidazole). Elution fractions containing the protein were pooled and concentrated (Amicon Ultra 100 kDa cut off, Millipore Sigma) for gel filtration. Gel filtration was conducted using a Superose 6 10/300 GL column (Cytiva) pre-equilibrated with GF buffer (20 mM Tris pH 8.0, 150 mM NaCl). Peak fractions corresponding to soluble protein were pooled and concentrated to 4.5–5.5 mg/mL (Amicon Ultra 100 kDa cut off, Millipore Sigma). Protein samples were flash-frozen in liquid nitrogen and stored at −80°C.

For purification of the RBD and NTD constructs, supernatant was harvested after 7 days of expression and incubated with 300 μL of Ni-NTA resin (Qiagen) at 4°C overnight. The resin was washed three times with 5 mL of PBS, then three times with 5 mL of PBS supplemented with 20 mM of imidazole. Proteins were eluted in PBS containing 300 mM of imidazole and then buffer exchanged into PBS and concentrated to 5-10 mg/mL (Amicon Ultra 10 kDa cut off, Millipore Sigma) before flash freezing and storage at −80°C.

### Antibody Production

VH-FC ab8, IgG ab1, Fab S309, and Fab S2M11 were produced as previously described^4,13,14^. Plasmids encoding light and heavy chains for Fab 4A8 and Fab 4-8 were synthesized (GeneArt Gene Synthesis, Thermo Fischer Scientific). Heavy chains were designed to incorporate a C terminal 6x histidine tag. Expi293 cells were transfected at a density of 3 x 10^6 cells/mL using linear polyethylenimine (Polysciences Cat# 23966-1). 24-hours following transfection, media was supplemented with 2.2 mM valproic acid, and expression was carried out for 3–5 days at 37°C, 8% CO_2_. The supernatant was harvested by centrifugation and filtered through a 0.22 μM filter prior to loading onto a 5 mL HisTrap excel column (Cytiva). The column was washed for 20 CVs with wash buffer (20 mM Tris pH 8.0, 500 mM NaCl), 5 CVs of wash buffer supplemented with 20 mM imidazole. The protein was eluted with elution buffer (20 mM Tris pH 8.0, 500 mM NaCl, 500 mM imidazole). Elution fractions containing the protein were pooled and concentrated (Amicon Ultra 10 kDa cut off, Millipore Sigma) for gel filtration. Gel filtration was conducted using a Superose 6 10/300 GL column (Cytiva) pre-equilibrated with GF buffer (20 mM Tris pH 8.0, 150 mM NaCl). Peak fractions corresponding to soluble protein were pooled and concentrated to 8–20 mg/mL (Amicon Ultra 10 kDa cut off, Millipore Sigma). Protein samples were stored at 4°C until use.

### Electron Microscopy Sample Preparation and Data Collection

For cryo-EM, S protein samples were prepared at 2.25 mg/mL, with and without the addition of ACE2 or antibody (1:1.25 S protein trimer:ACE2 molar ratio, 1:3 S protein trimer:antibody molar ratio). Vitrified samples of all S protein samples were prepared by first glow discharging Quantifoil R1.2/1.3 Cu mesh 200 holey carbon grids for 30 seconds using a Pelco easiGlow glow discharge unit (Ted Pella) and then applying 1.8 μL of protein suspension to the surface of the grid at a temperature of 10°C and a humidity level of >98%. Grids were blotted (12 sec, blot force −10) and plunge frozen into liquid ethane using a Vitrobot Mark IV (Thermo Fisher Scientific). All cryo-EM samples were imaged using a 300 kV Titan Krios G4 transmission electron microscope (Thermo Fisher Scientific) equipped with a Falcon4 direct electron detector in electron event registration (EER) mode. Movies were collected at 155,000x magnification (calibrated pixel size of 0.5 Å per physical pixel) over a defocus range of −0.5 μm to −3 μm with a total dose of 40 e^-^/Å^2^ using EPU automated acquisition software.

### Image Processing

The detailed workflow for the data processing is summarized in Supplementary Figures S1-S4, S6-S7, S11-S14, S17, S23, S25. In general, all data processing was performed in cryoSPARC v.3.2^36^ unless stated otherwise. Patch mode motion correction (EER upsampling factor 1, EER number of fractions 40), patch mode CTF estimation, reference free particle picking, and particle extraction were carried out on-the-fly in cryoSPARC live. After preprocessing, particles were subjected to 2D classification and/or 3D heterogeneous classification. The final 3D refinement was performed with estimation of per particle CTF and correction for high-order aberrations.

For the complexes of spike protein ectodomain and human ACE2, focused refinements were performed with a soft mask covering a single RBD and its bound ACE2. For the complexes of spike protein ectodomain and V_H_ ab6, a soft mask covering V_H_-ab6 and its bound RBD was used in focused refinement. For the complexes of Gamma spike protein ectodomain and fab 4-8/4-A8, another round of 3D refinement with C3 symmetry was carried out, followed by symmetry expansion. The derived particles were then focused-refined with a soft mask covering single NTD and its bound fab.

Global resolution and focused resolution were determined according to the gold-standard FSC^37^.

### Model Building and Refinement

Initial models either from published coordinates (PDB code 7MJG, 7MJM, 7MJN, 7LXY, 7K43 and 7MJI) or from homology modeling (for 4-8, 4-A8 and V_H_-ab6) were docked into the focused refinement maps or global refinement maps using UCSF Chimera v.1.15^38^. Then, mutation and manual adjustment were performed with COOT v.0.9.3^39^, followed by iterative rounds of refinement in COOT and Phenix v.1.19^40^. Model validation was performed using MolProbity^41^. Figures were prepared using UCSF Chimera, UCSF ChimeraX v.1.1.1^42^, and PyMOL (v.2.2 Schrodinger, LLC).

### Biolayer Interferometry (BLI) S protein - ACE2 Binding Assay

The kinetics of SARS-CoV-2 trimers and human ACE2 binding were analyzed with the biolayer interferometer BLItz (ForteBio, Menlo Park, CA). Protein-A biosensors (ForteBio: 18–5010) were coated with ACE2-mFc (40 μg/mL) for 2 min and incubated in DPBS (pH = 7.4) to establish baselines. Concentrations of 125, 250, 500, and 1000 nM spike trimers were used for association for 2 min followed by dissociation in DPBS for 5 min. The association (*k*_on_) and dissociation (*k*_off_) rates were derived from the fitting of sensorgrams and used to calculate the binding equilibrium constant (K_D_).

### Flow Cytometry S protein - ACE2 Binding Assay

Expi293 cells were transfected with plasmids encoding the various full length variant S proteins using linear PEI as performed during recombinant spike production. After 3 days of expression, cells were washed with PBS then pelleted and resuspended in incubation buffer (PBS pH 7.4 with 0.02% Tween-20 and 4% BSA) at a concentration of 5 x 10^6 cells/ml. 100 microliters of this mixture was deposited into each well within a 96 well plate followed by a 5 minute incubation on ice. Cells were pelleted and resuspended in incubation buffer containing increasing concentrations of recombinant FC-ACE2 (Sino Biological Cat# 10108-H05H-100) followed by incubation for 20 minutes on ice. The cells were pelleted and washed with 200 microliters of wash buffer (PBS pH 7.4 with 0.02%Tween-20). Next, the cells were incubated in 100 microliters of a 1:300 dilution of secondary antibody (Thermo Fisher Cat# A-21235) in incubation buffer for 10 minutes. Cells were pelleted and washed twice in 100 microliters of wash buffer prior to staining with propidium iodide (Biolegend). Cells were analyzed using a Cytoflex LX at the ubcFLOW cytometry facility. Data was analyzed using FlowJo. The gating strategy employed along with results for un-transfected (negative control) Expi293 cells are available in figure S10B.

### Enzyme-Linked Immunosorbent Assay (ELISA)

For ELISA, 100 μL of wild-type (D614G), or variant SARS-CoV-2 S proteins, NTDs, and RBDs were coated onto 96-well MaxiSorp^™^ plates at 2 μg/mL in PBS + 1% casein overnight at 4°C. All washing steps were performed 3 times with PBS + 0.05% Tween 20 (PBS-T). After washing, wells were incubated with blocking buffer (PBS-T + 1% casein) for 1 hr at room temperature. After washing, wells were incubated with dilutions of primary antibodies in PBS-T + 0.5% BSA buffer for 1 hr at room temperature. After washing, wells were incubated with goat anti-human IgG (Jackson ImmunoResearch) at a 1:8,000 dilution in PBS-T + 1% casein buffer for 1 hr at room temperature. After washing, the substrate solution (Pierce^™^ 1-Step^™^) was used for colour development according to the manufacturer’s specifications. Optical density at 450 nm was read on a Varioskan Lux plate reader (Thermo Fisher Scientific).

### Authentic SARS-CoV-2 Plaque Reduction Neutralization Assay

Neutralization assays were performed using Vero E6 cells (ATCC CRL-1586) that were seeded 24 hours prior to the assay in 24-well tissue culture plates at a density of 3 x 10^5 cells per well. Antibodies were serially diluted two-fold (starting concentration of 4 ug/mL, 10 ug/mL, or 40 ug/mL, depending on the antibody being tested) and mixed with equal volume of 30-50 plaque forming units of SARS-CoV-2. This results in a final antibody concentration of 2 ug/mL, 5 ug/mL, or 20 ug/mL in the antibody-virus mixture. The following SARS-CoV-2 variants were used: isolate USA-WA1/2020 (NR-52281, BEI Resources); isolate hCoV-19/South Africa/KRISP-EC-K005321/2020 (NR-54008, BEI Resources); isolate hCoV-19/USA/CA_UCSD_5574/2020 (NR-54008, BEI Resources); and isolate hCoV-19/USA/PHC658/2021 (NR-55611, BEI Resources). The antibody-virus mixture was then incubated at 37°C in a 5% CO_2_ incubator for 1 hour and added to the Vero E6 cell seeded monolayers, in duplicate. Plates were then incubated for 1 hour at 37°C in a 5% CO_2_ incubator. Following incubation, an overlay media with 1% agarose-containing media (2x Minimal Essential Medium, 7.5% bovine albumin serum, 10 mM HEPES, 100 μg/mL penicillin G and 100 U/mL streptomycin) was added to the monolayers. The plates were incubated for 48-72 hours (depending on the SARS-CoV2 variant) and then cells were fixed with formaldehyde for 2 hours. Following fixation, agar plugs were removed, and cells were stained with crystal violet. In order to assess the input virus, a viral back-titration was performed using culture medium as a replacement for the antibodies. To estimate the neutralizing capability of each antibody, IC50s were calculated by non-linear regression using the sigmoidal dose response equation in GraphPad Prism 9. All assays were performed in the University of Pittsburgh Regional Biocontainment Laboratory BSL-3 facility.

