## Supplemental Information for "Emerging SARS-CoV-2 Variants of Concern: Spike Protein Mutational Analysis and Epitope for Broad Neutralization"

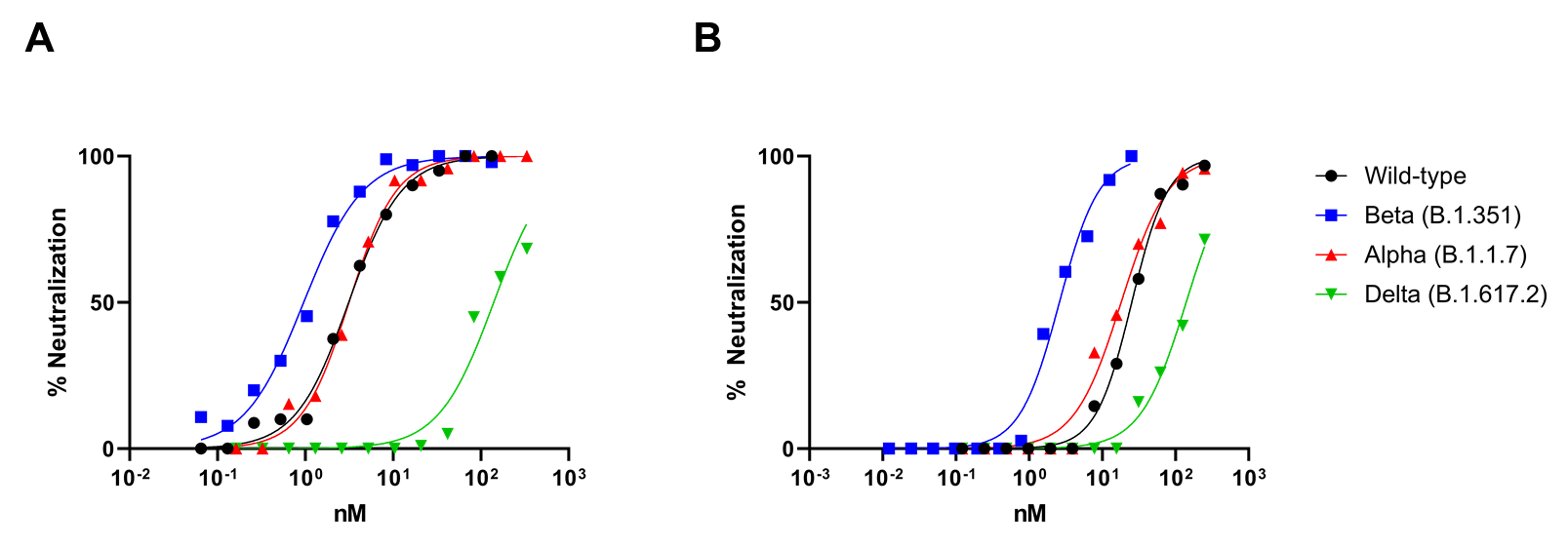

**Figure S1. Live virus neutralization assays using ab6.** Either V_H_ ab6 **(A)** or V_H_-FC ab6 **(B)** was used in the plaque reduction neutralization test - PRNT. Experiments were performed in duplicate, and the means are plotted.

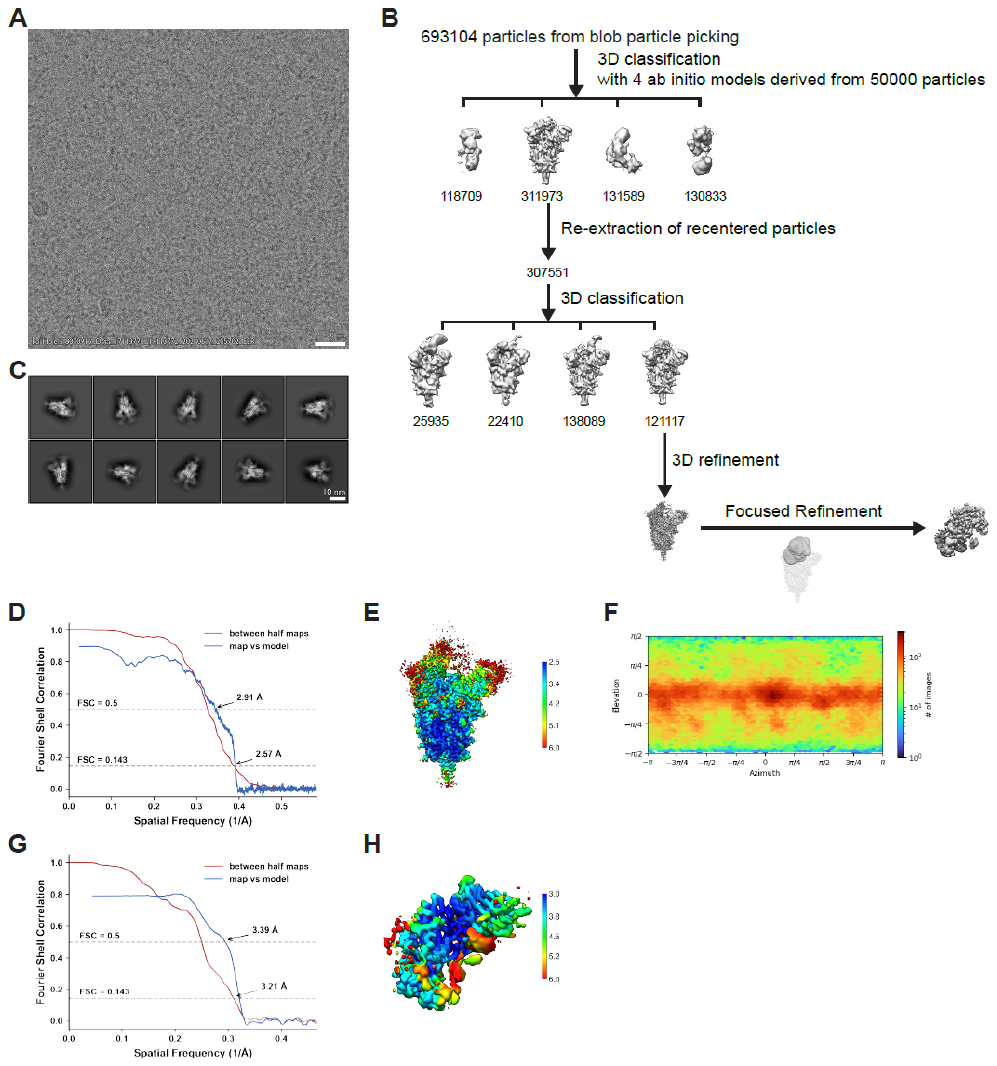

**Figure S2. Cryo-EM data processing and validation for complex of D614G spike protein ectodomain and V_H_-ab6.** **(A)** Representative cryo-EM micrograph. **(B)** Workflow of cryo-EM image processing. **(C)** Representative 2D classes. **(D-F)** FSC curves (D), local resolution (E) and viewing direction distribution plot (F) of global refinement. **(G-H)** FSC curves (G) and local resolution (H) of focused refinement.

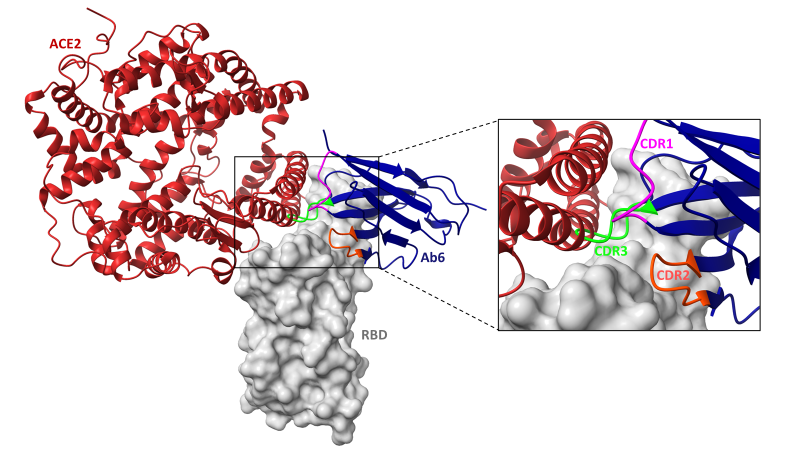

**Figure S3. Overlap of ab6 and ACE2 footprints.** The local refined model of the ab6-RBD interface was superposed with the crystal structure of the ACE2-RBD complex (PDB: 6M0J). ACE2 is shown in red cartoon while V_H_ ab6 is shown in blue cartoon, with each CDR loop coloured. The RBD is depicted as a grey molecular surface. Models were aligned using the RBD for superposition

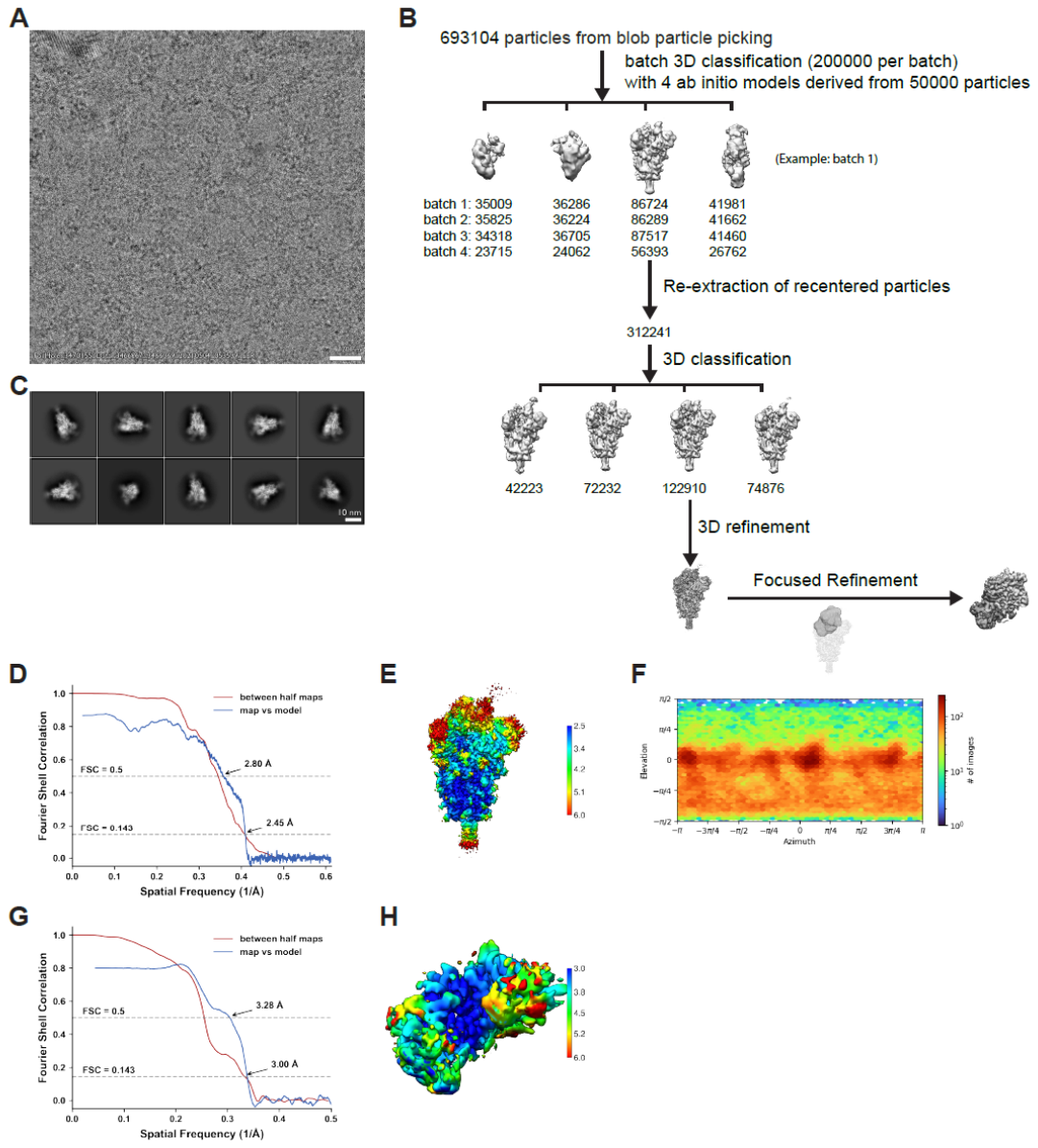

**Figure S4. Cryo-EM data processing and validation for complex of Epsilon spike protein ectodomain and V_H_-ab6.** **(A)** Representative cryo-EM micrograph. **(B)** Workflow of cryo-EM image processing. **(C)** Representative 2D classes. **(D-F)** FSC curves (D), local resolution (E) and viewing direction distribution plot (F) of global refinement. **(G-H)** FSC curves (G) and local resolution (H) of focused refinement.

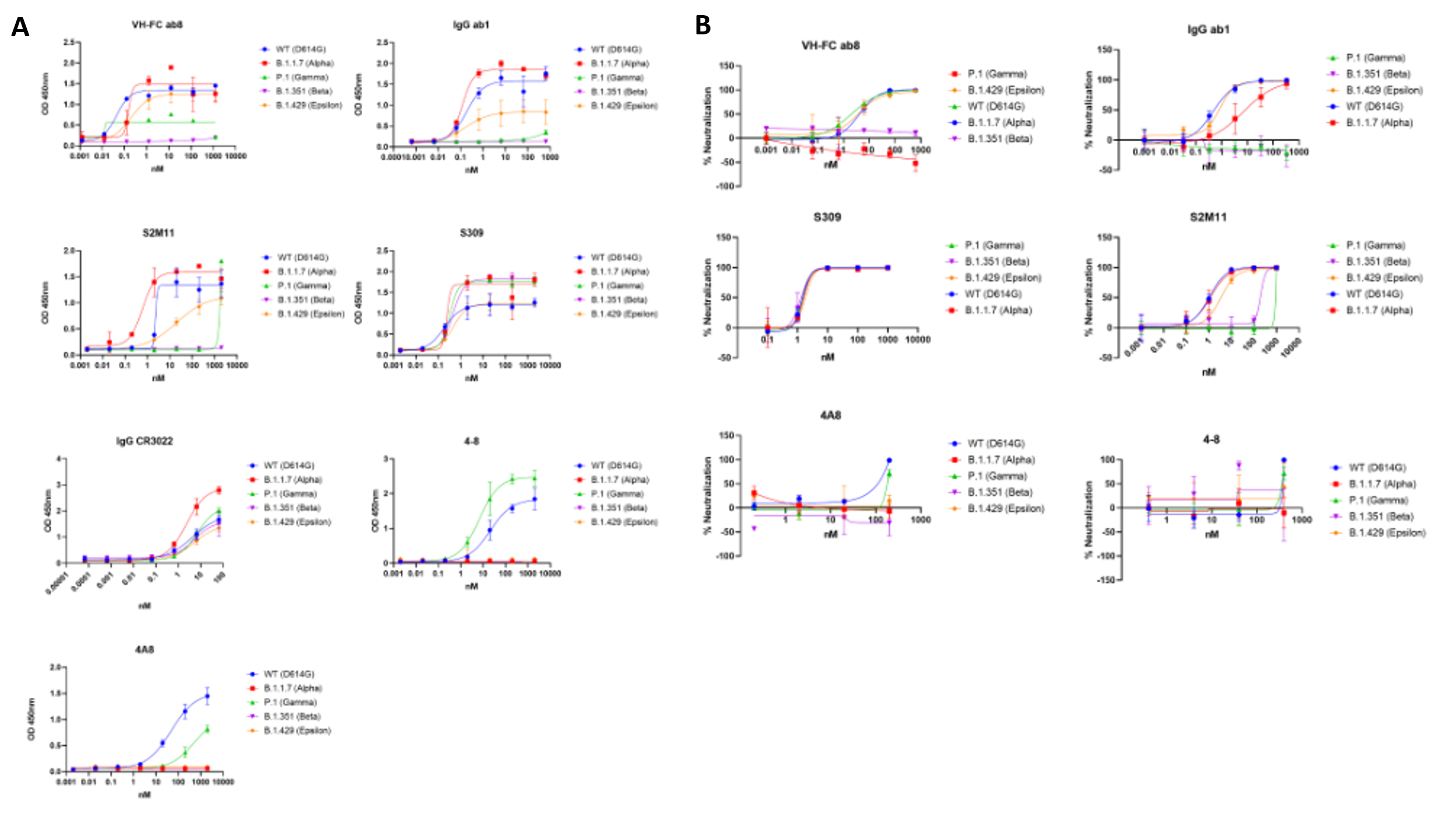

**Figure S5. Antibody binding and neutralization curves.** Related to Figure 2B. **(A)** Antibody binding curves as determined via ELISA. **(B)** Antibody neutralization of pseudoviruses. Experiments were performed in triplicate. Error bars denote the standard error of the mean.

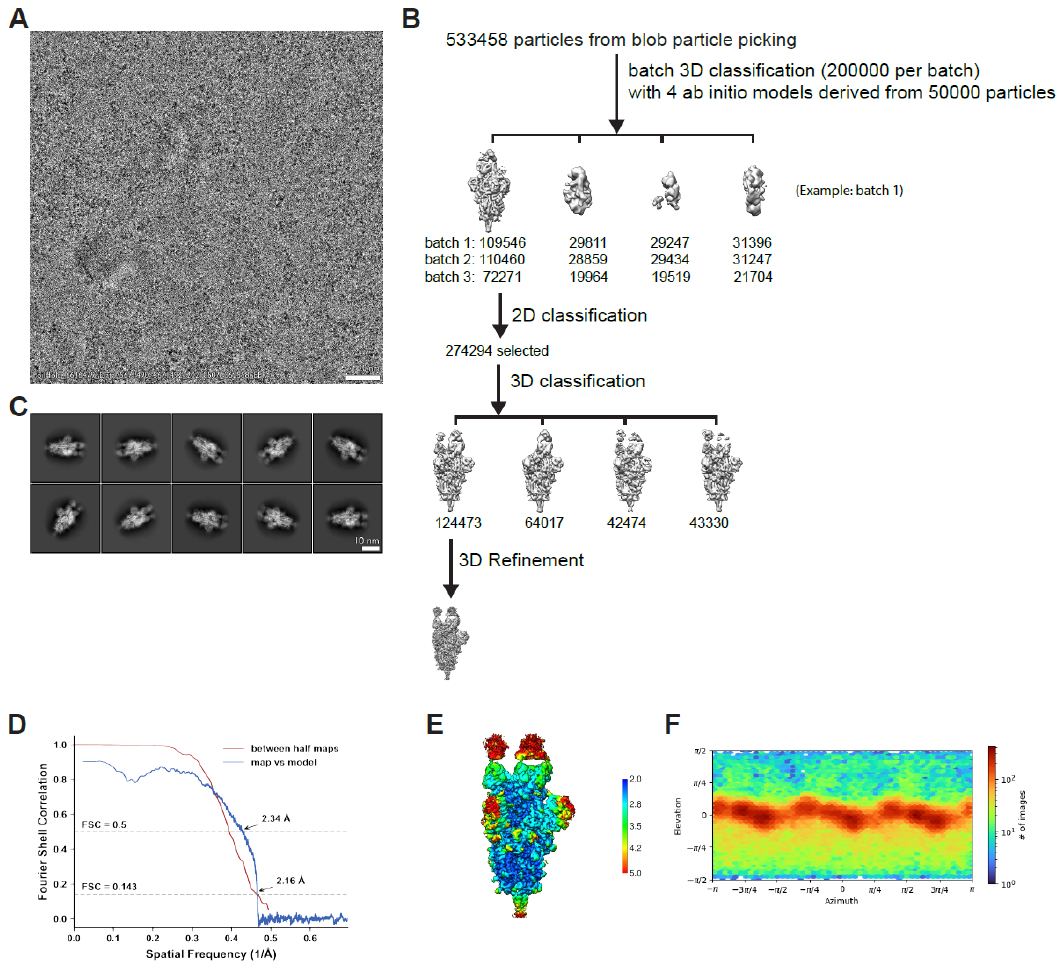

**Figure S6. Cryo-EM data processing and validation for complex of Epsilon spike protein ectodomain and S2M11.** **(A)** Representative cryo-EM micrograph. **(B)** Workflow of cryo-EM image processing. **(C)** Representative 2D classes. **(D)** FSC curves. **(E)** Local resolution. **(F)** Viewing direction distribution plot.

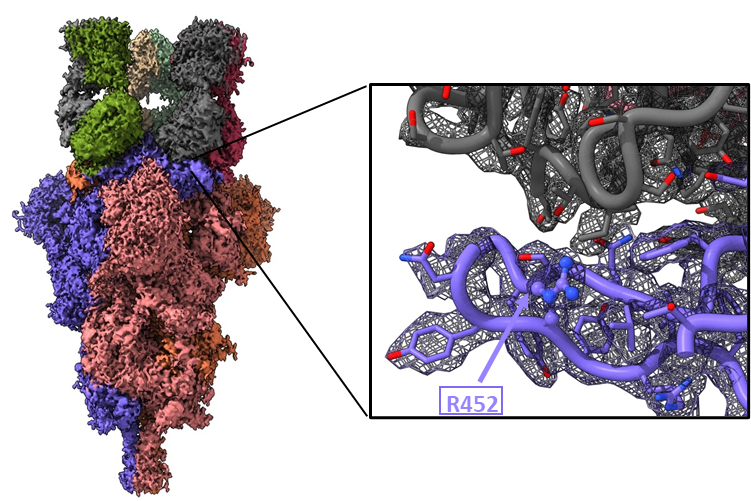

**Figure S7. Structure of the Epsilon Variant Spike – S2M11 complex.** Global map of the Epsilon Variant Spike – S2M11 complex and zoomed in view of map and model at the S2M11(Grey)-RBD(Purple) interface. R452 is indicated with an arrow and boxed label.

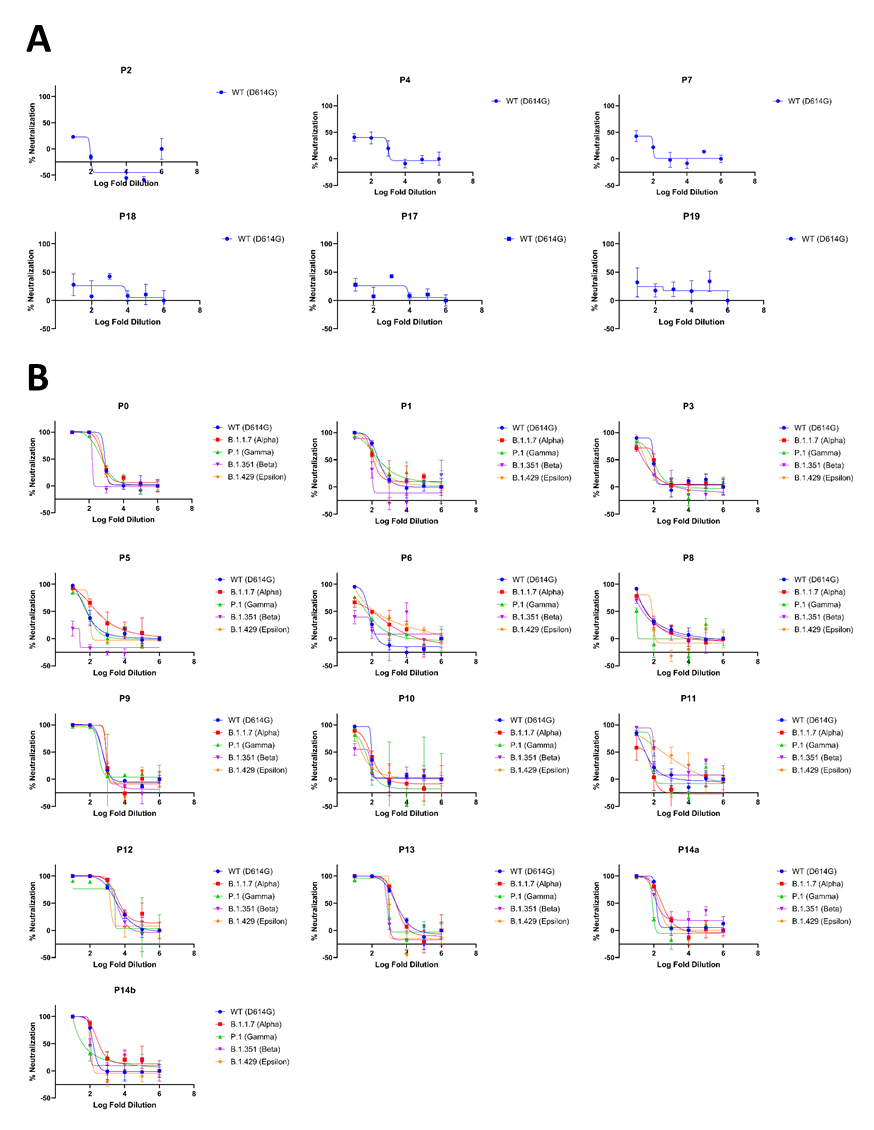

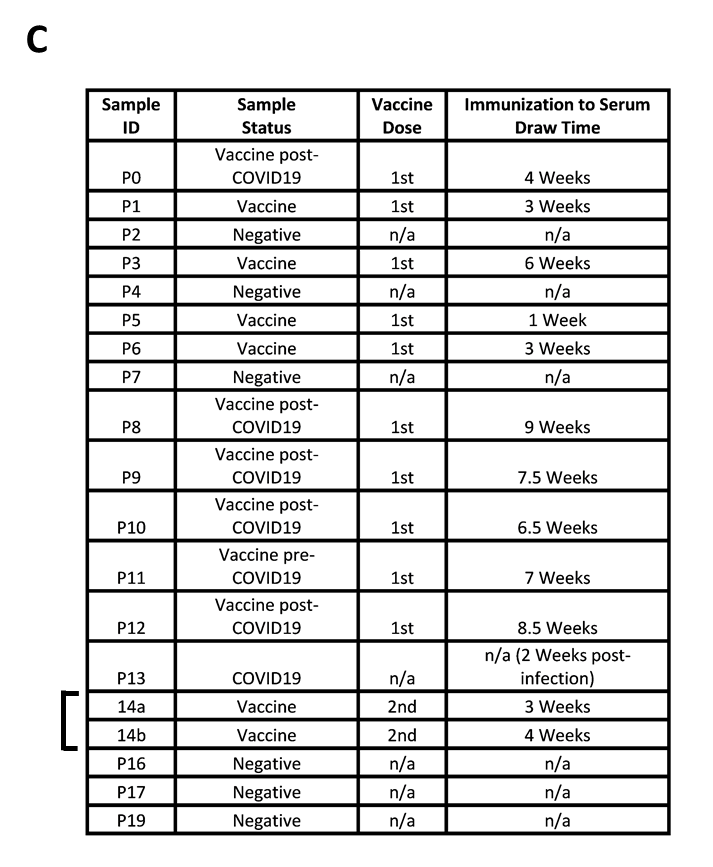

**Figure S8. Patient-derived sera sample information and raw pseudovirus neutralization data.** Related to figure 2C-D. **(A)** Neutralization of wild type spike pseudotyped virus using pre-pandemic sera samples. **(B)** Neutralization of pseudoviruses by the indicated patient sera. **(C)** Patient-derived sera sample information including sample number, vaccination status, vaccination dose number and immunization to serum draw time. (n/a: not applicable). Experiments were performed in triplicate. Error bars denote the standard error of the mean.

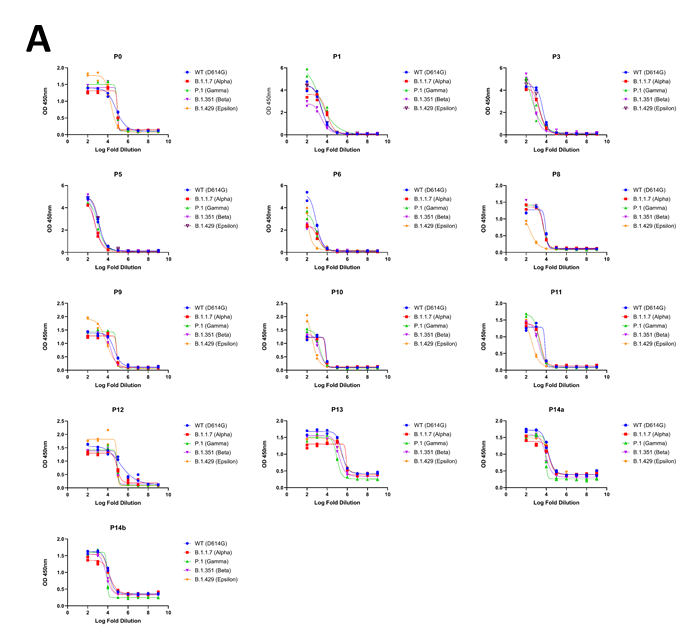

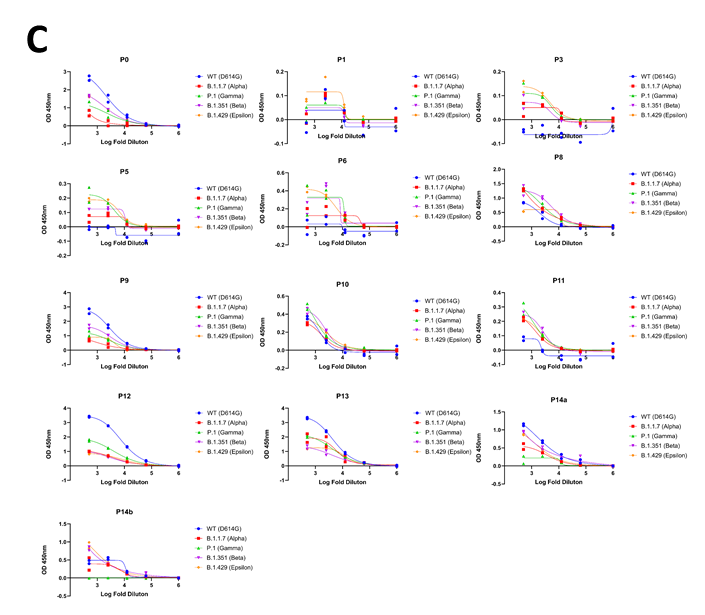

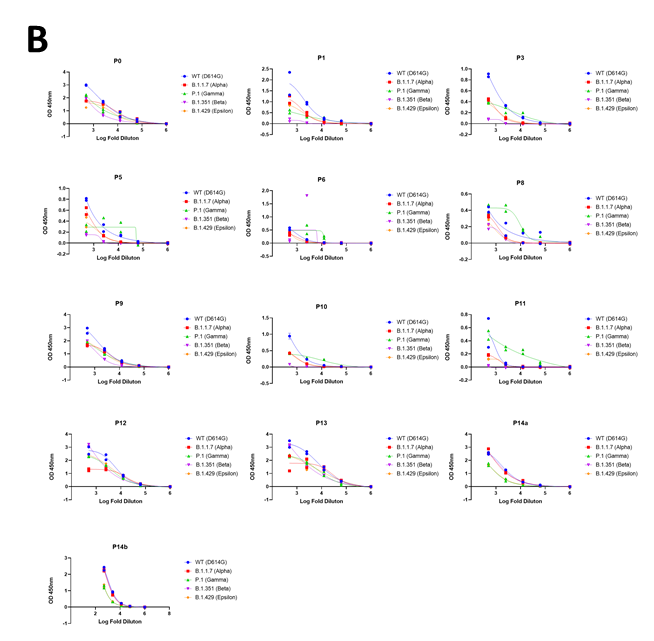

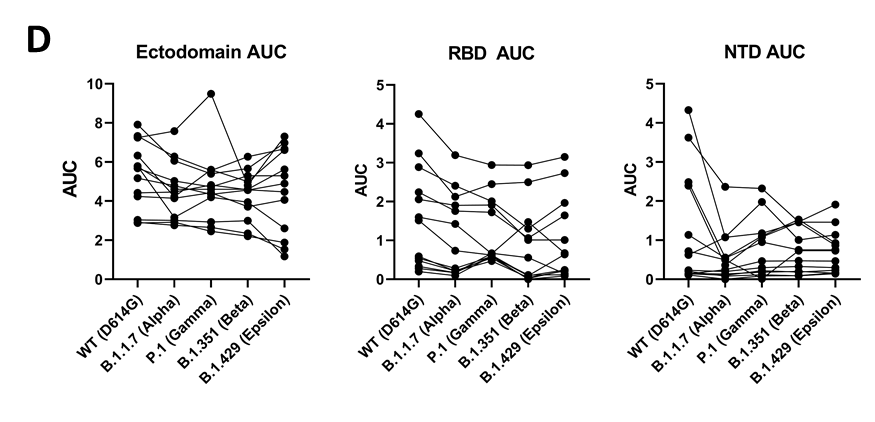

**Figure S9. Ectodomain, RBD, and NTD binding by antibodies in patient sera.** Related to Figure 2D. Binding of wild type or variant ectodomains **(A)**, RBDs **(B)**, or NTDs **(C)** as assessed by ELISA. Experiments were performed in duplicate, and the results are plotted as points. **(D)** Aggregated area under the curve (AUC) values for each serum sample, protein construct, and spike variant.

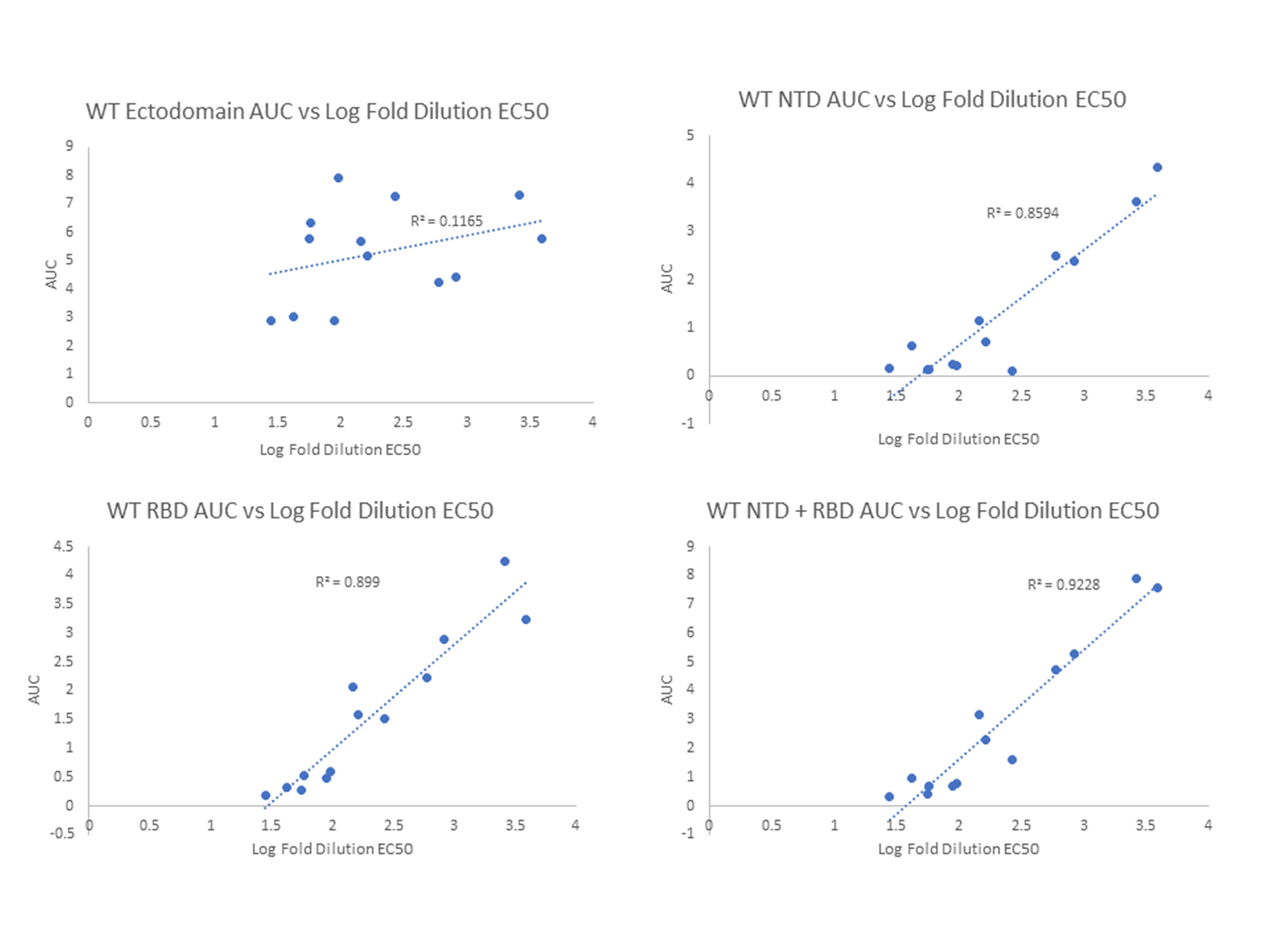

**Figure S10. High correlation between NTD and RBD binding antibody levels and pseudoviral neutralization in patient derived sera.** Wild type (WT) Ectodomain, NTD, and RBD binding by patient sera was assessed via ELISA and area under the curve (AUC) was calculated from the resulting data. AUC’s were correlated with neutralization potencies for wild type spike pseudo typed virus via linear regression. Correlation coefficients are shown for each comparison.

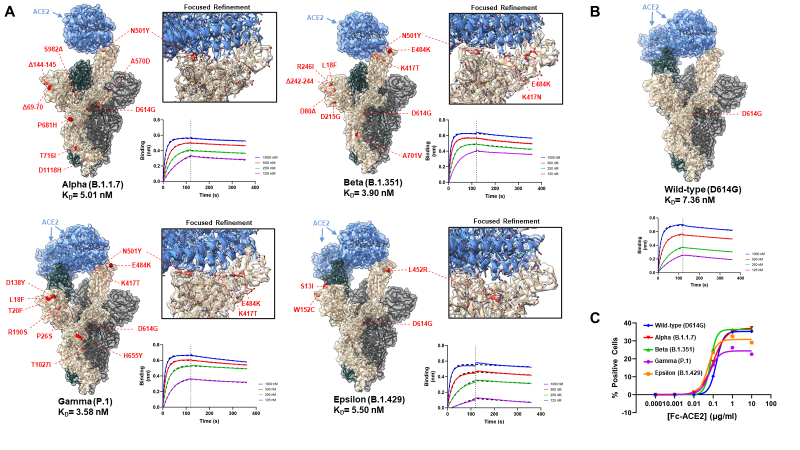

**Figure S11. Structural and Functional Analysis of ACE2 Binding by Variant Spikes. (A)** CryoEM and biolayer inferometry (BLI) analysis of ACE2 binding by variant spikes. Shown for each variant are global spike-ACE2 complex models with mutational positions highlighted as red spheres (locations of mutations which can not be modelled are approximated as occurring at the nearest modelled residue), maps and models of the ACE2-RBD interface obtained via focused refinement, and BLI sensorgrams for ACE2-spike binding experiments. **(B)** Global spike-ACE2 complex model and BLI sensorgram for the wild-type spike. **(C)** ACE2 binding of cells expressing full-length spikes was measured by Flow-cytometry.

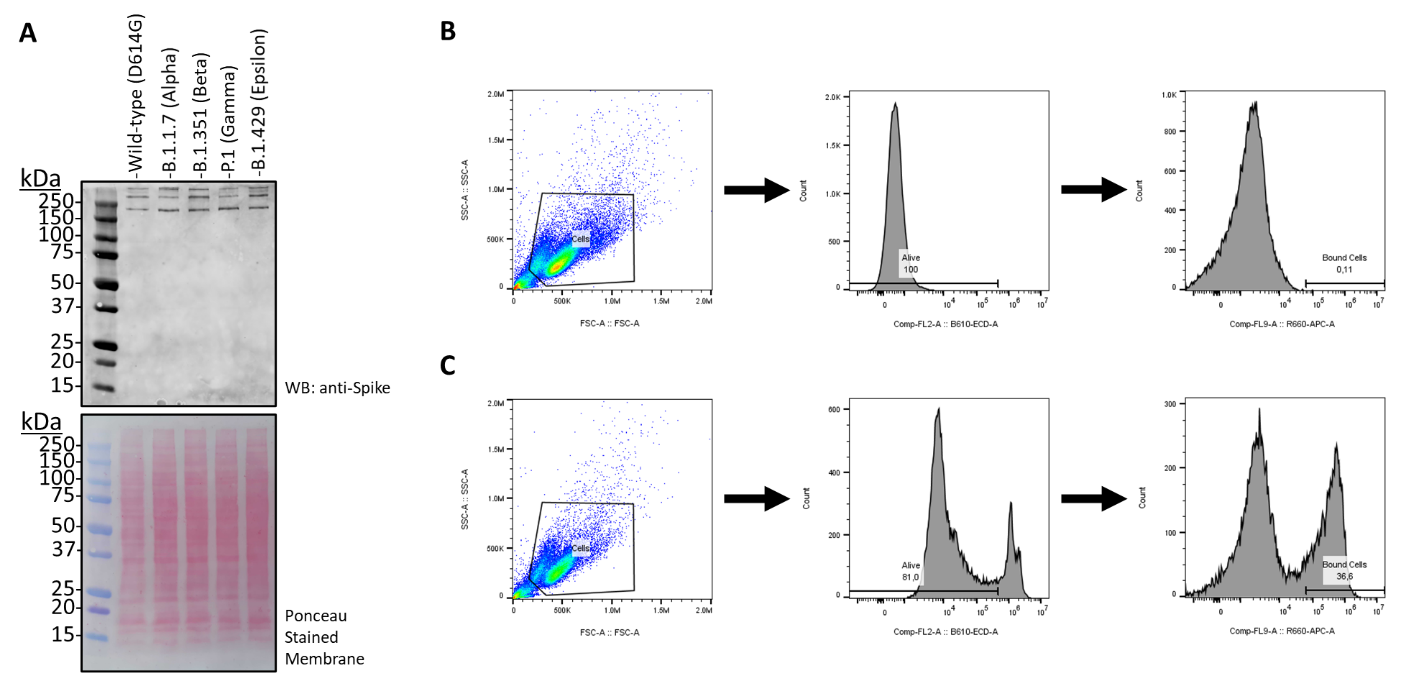

**Figure S12. Full length spike protein expression and flow cytometry gating strategy.** **(A)** Western blot of full-length spike proteins expressed in Expi293 cells. The ponceau stained membrane is shown as a loading control. **(B-C)** Gating strategy used for flow cytometry experiments. Panel (B) depicts signals obtained for un-transfected Expi293 cells incubated with the highest concentration of ACE2 utilized (10μg/ml) and panel (C) depicts signals obtained for spike expressing Expi293 cells under the same conditions.

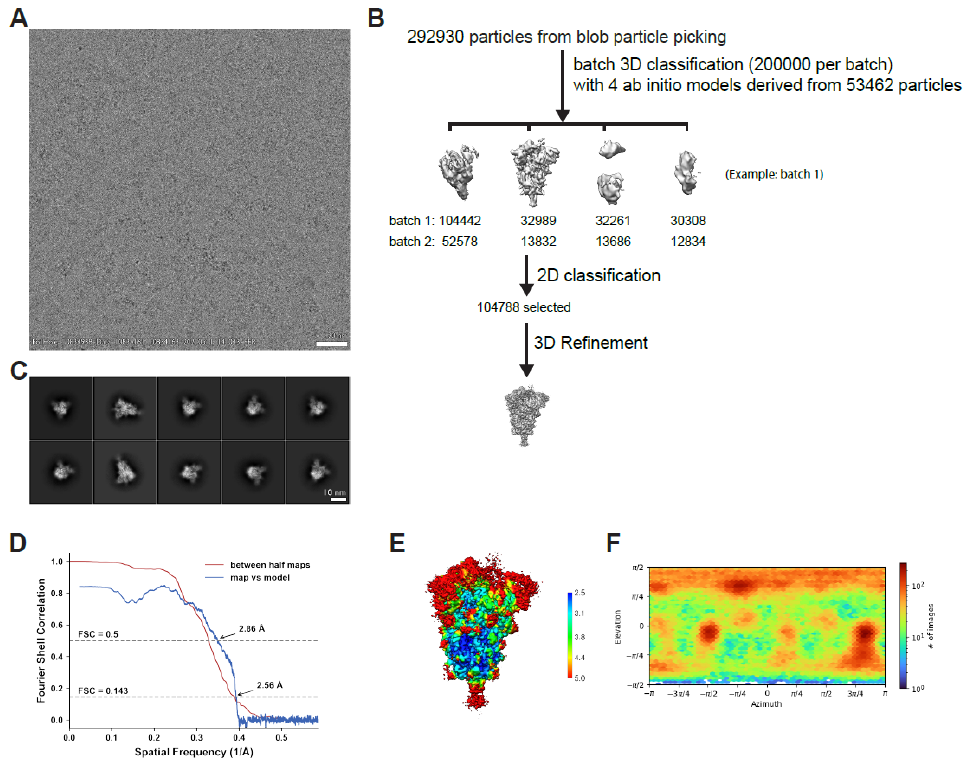

**Figure S13. Cryo-EM data processing and validation for the Alpha spike protein ectodomain.**

**(A)** Representative cryo-EM micrograph. **(B)** Workflow of cryo-EM image processing. **(C)** Representative 2D classes. **(D)** FSC curves. **(E)** Local resolution. **(F)** Viewing direction distribution plot.

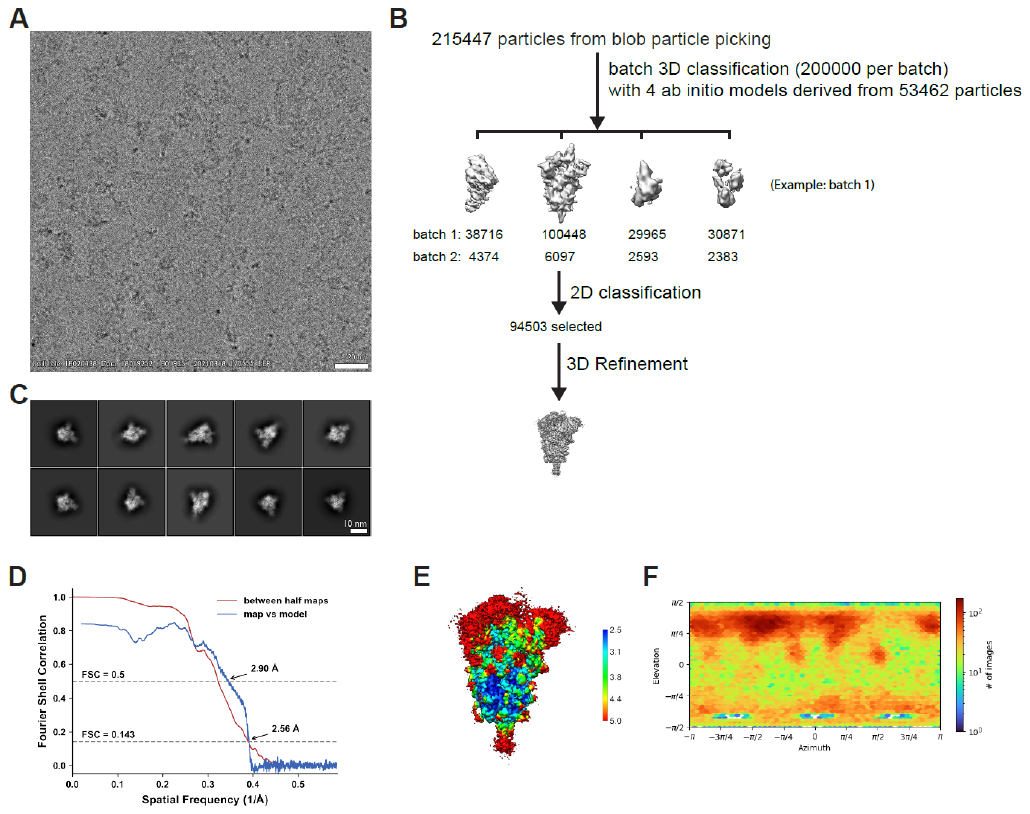

**Figure S14. Cryo-EM data processing and validation for the Beta spike protein ectodomain.**

**(A)** Representative cryo-EM micrograph. **(B)** Workflow of cryo-EM image processing. **(C)** Representative 2D classes. **(D)** FSC curves. **(E)** Local resolution. **(F)** Viewing direction distribution plot.

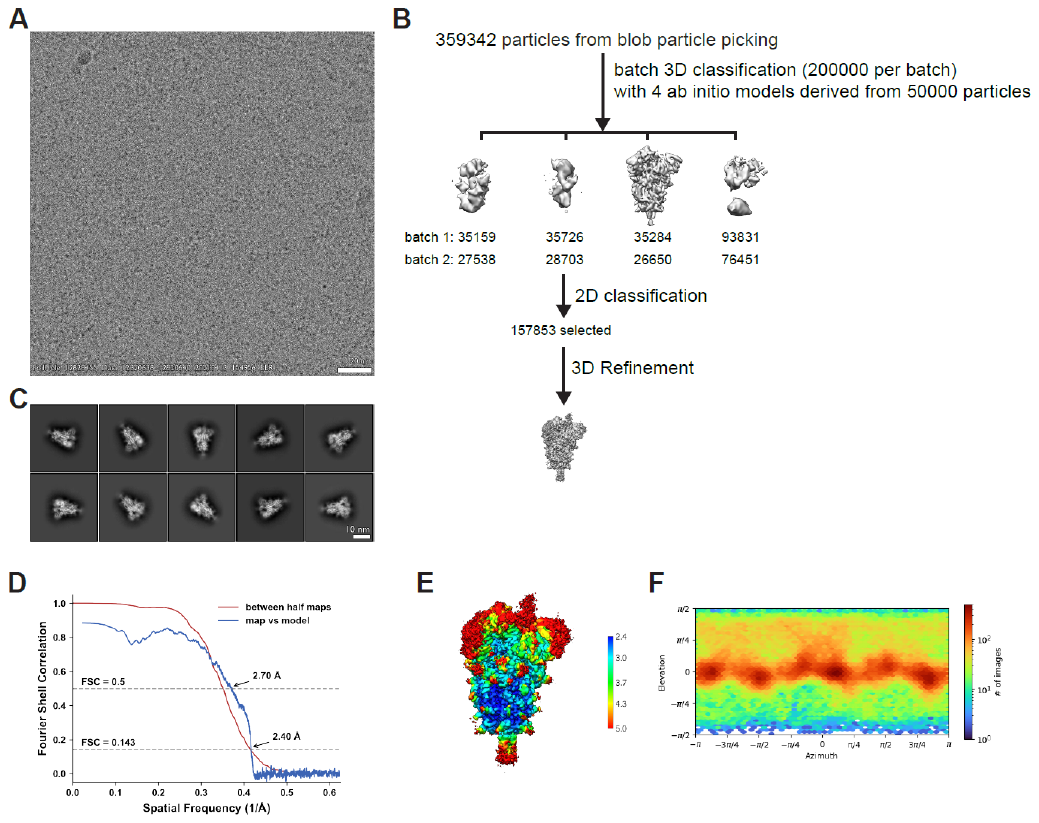

**Figure S15. Cryo-EM data processing and validation for the Epsilon spike protein ectodomain.**

**(A)** Representative cryo-EM micrograph. **(B)** Workflow of cryo-EM image processing. **(C)** Representative 2D classes. **(D)** FSC curves. **(E)** Local resolution. **(F)** Viewing direction distribution plot.

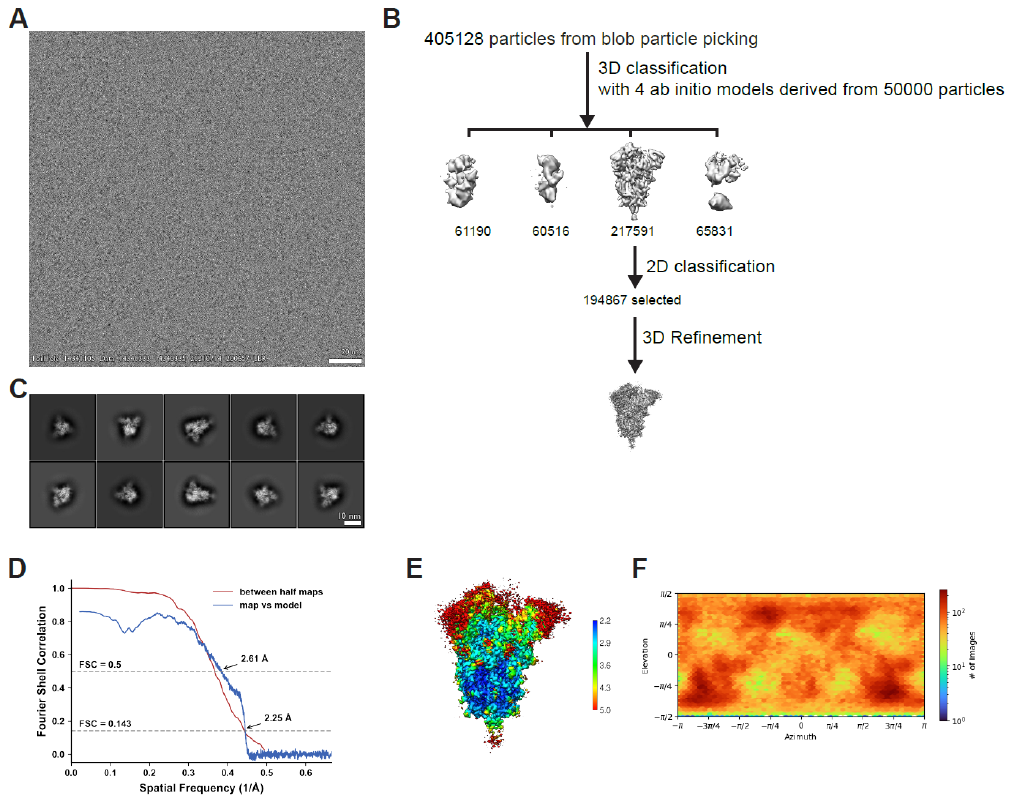

**Figure S16. Cryo-EM data processing and validation for the Gamma spike protein ectodomain.**

**(A)** Representative cryo-EM micrograph. **(B)** Workflow of cryo-EM image processing. **(C)** Representative 2D classes. **(D)** FSC curves. **(E)** Local resolution. **(F)** Viewing direction distribution plot.

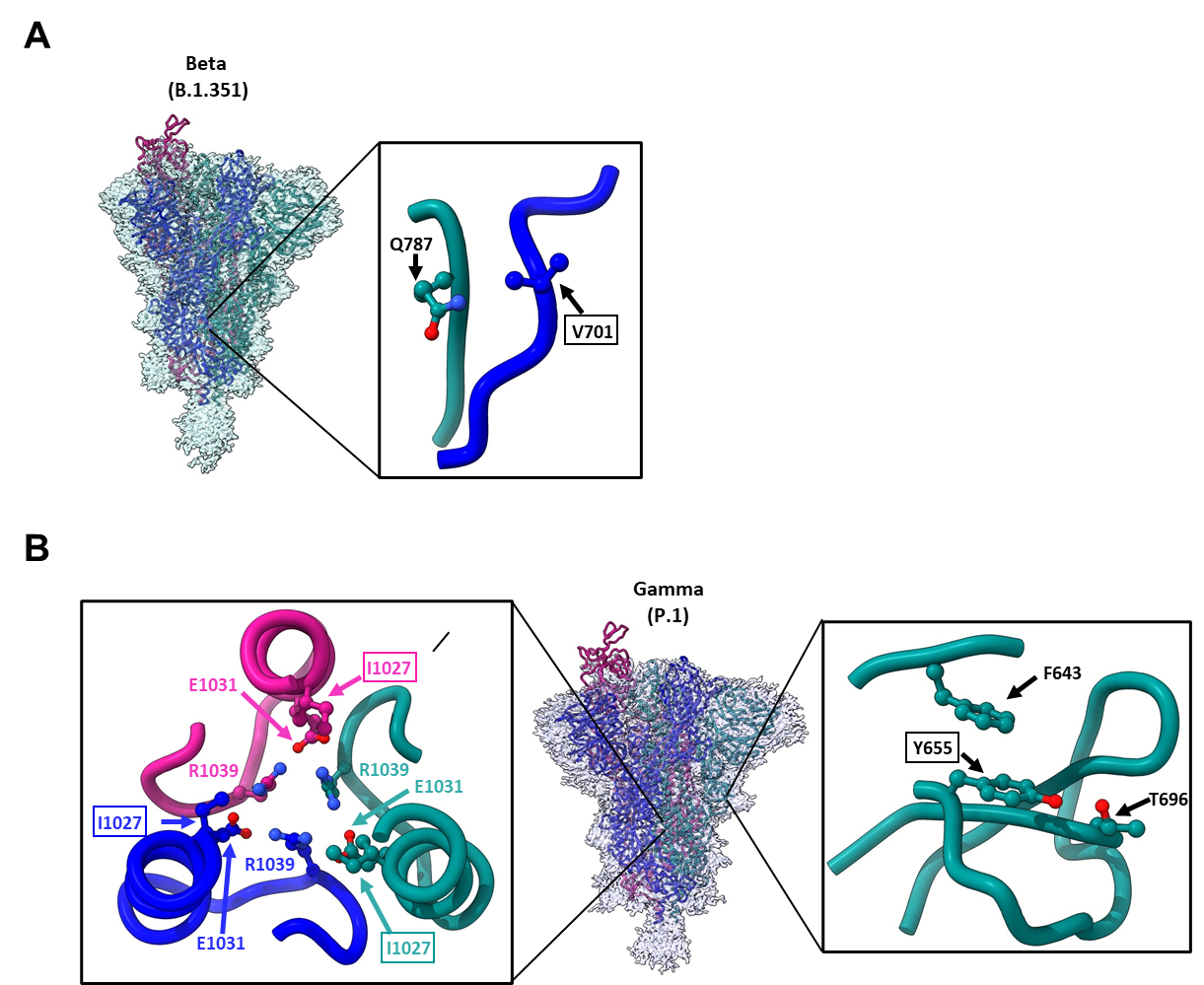

**Figure S17. Structural impacts of S2 mutations within the Beta and Gamma S protiens. (A)** Global map and model of the Beta variant spike along with a zoomed in view of the A701->V mutation. **(B)** Global map and model of the Gamma variant spike along with zoomed in views of the T1027->I and H655->Y mutations. Mutated residues are indicated as boxed labels.

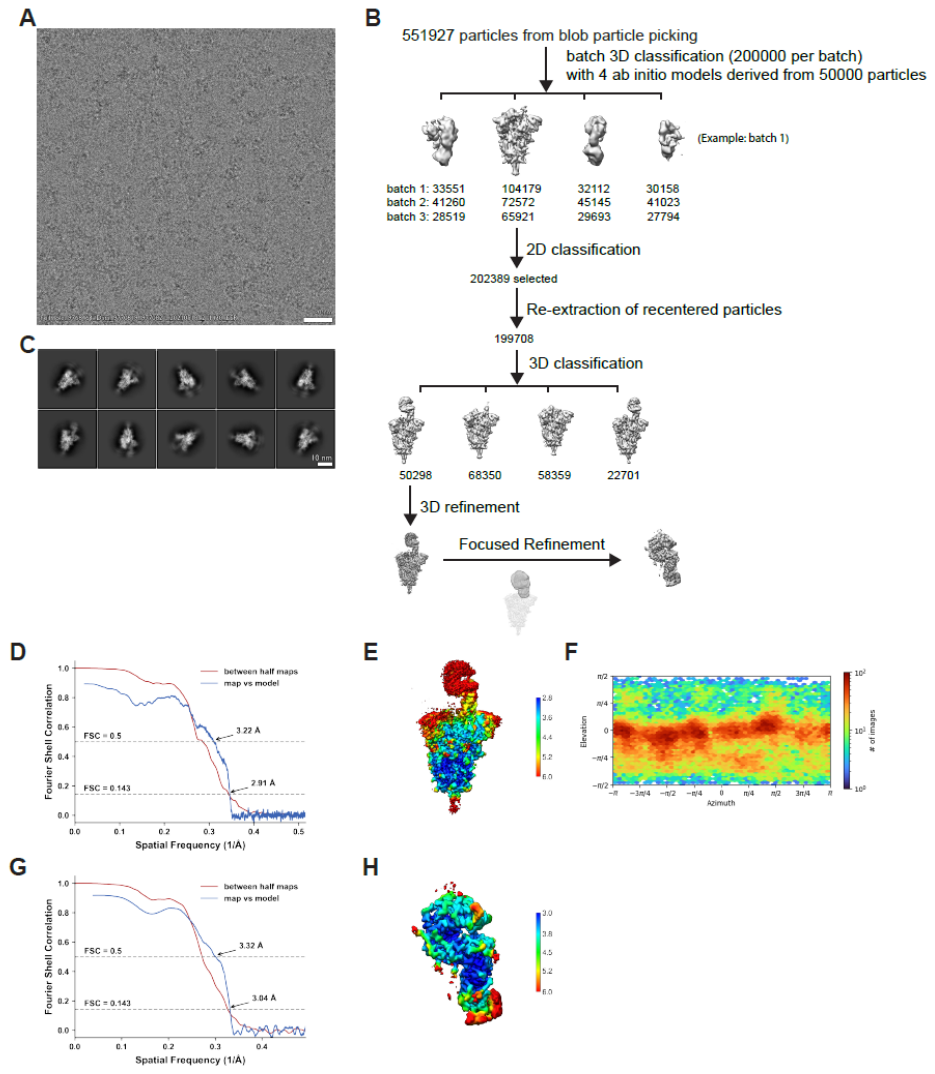

**Figure S18. Cryo-EM data processing and validation for complex of Alpha spike protein ectodomain and human ACE2.** **(A)** Representative cryo-EM micrograph. **(B)** Workflow of cryo-EM image processing. **(C)** Representative 2D classes. **(D-F)** FSC curves (D), local resolution (E) and viewing direction distribution plot (F) of global refinement. **(G-H)** FSC curves (G) and local resolution (H) of focused refinement.

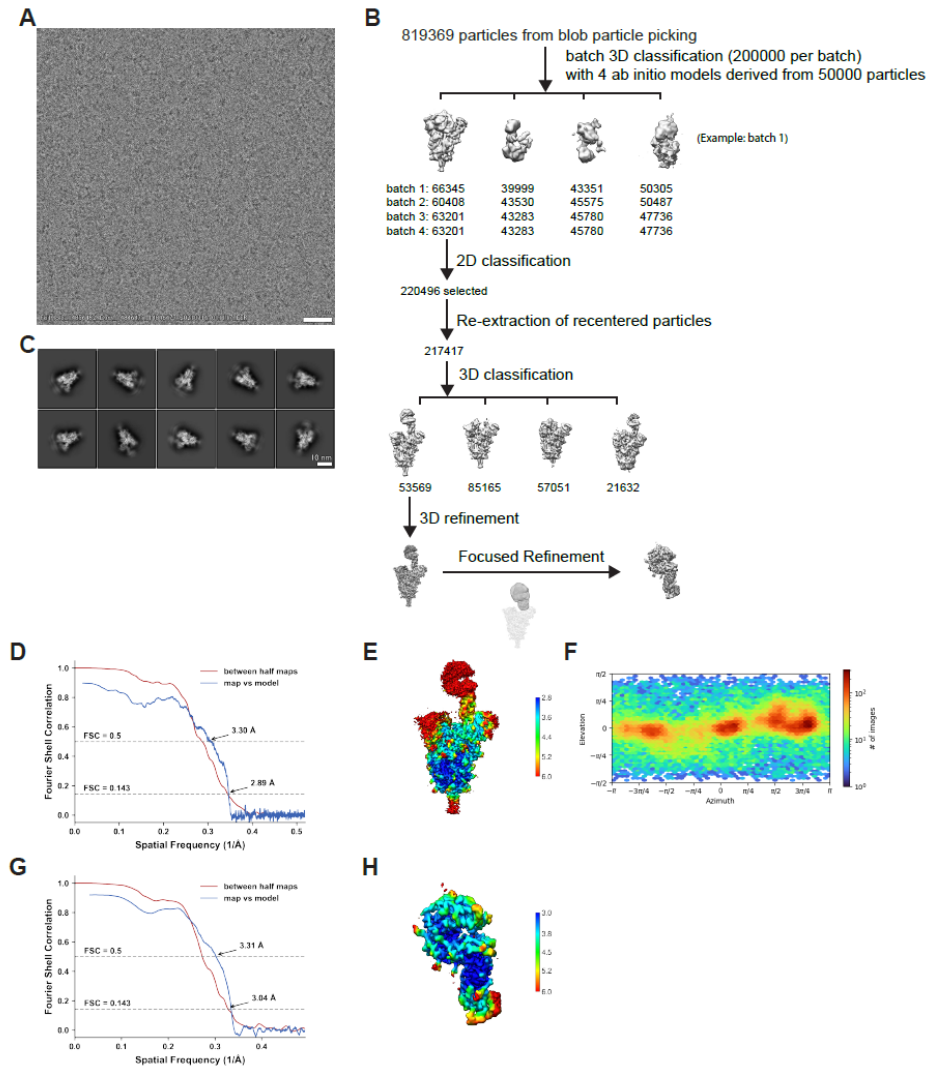

**Figure S19. Cryo-EM data processing and validation for complex of Beta spike protein ectodomain and human ACE2. (A)** Representative cryo-EM micrograph. **(B)** Workflow of cryo-EM image processing. **(C)** Representative 2D classes. **(D-F)** FSC curves (D), local resolution (E) and viewing direction distribution plot (F) of global refinement. **(G-H)** FSC curves (G) and local resolution (H) of focused refinement.

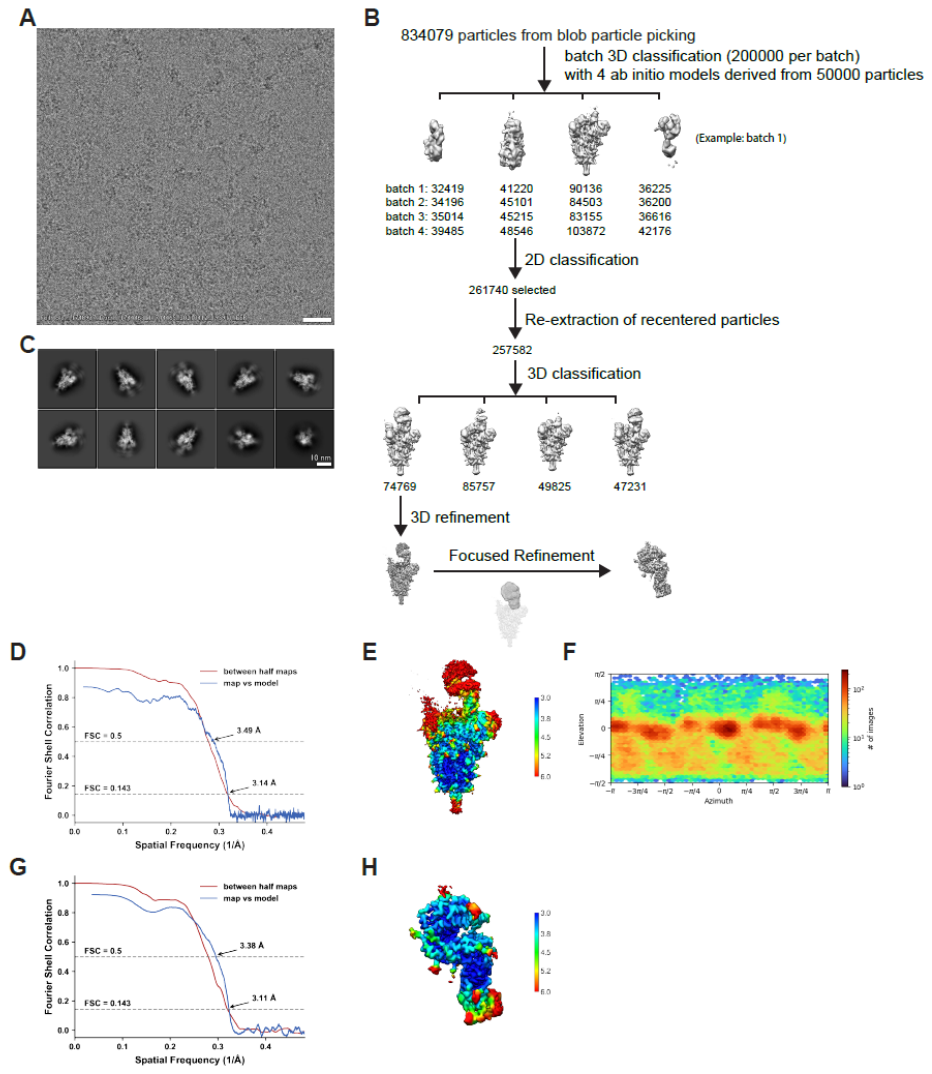

**Figure S20. Cryo-EM data processing and validation for complex of Epsilon spike protein ectodomain and human ACE2.** **(A)** Representative cryo-EM micrograph. **(B)** Workflow of cryo-EM image processing. **(C)** Representative 2D classes. **(D-F)** FSC curves (D), local resolution (E) and viewing direction distribution plot (F) of global refinement. **(G-H)** FSC curves (G) and local resolution (H) of focused refinement.

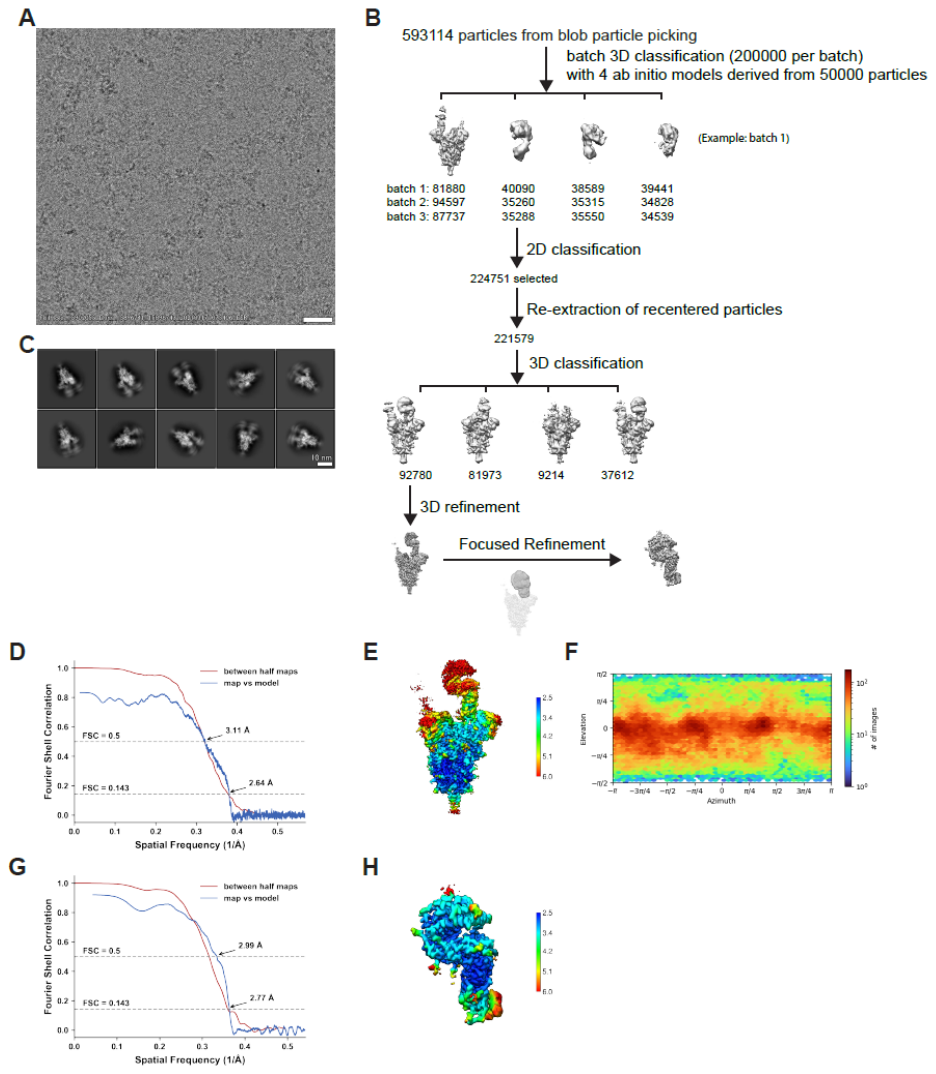

**Figure S21. Cryo-EM data processing and validation for complex of Gamma spike protein ectodomain and human ACE2. (A)** Representative cryo-EM micrograph. **(B)** Workflow of cryo-EM image processing. **(C)** Representative 2D classes. **(D-F)** FSC curves (D), local resolution (E) and viewing direction distribution plot (F) of global refinement. **(G-H)** FSC curves (G) and local resolution (H) of focused refinement.

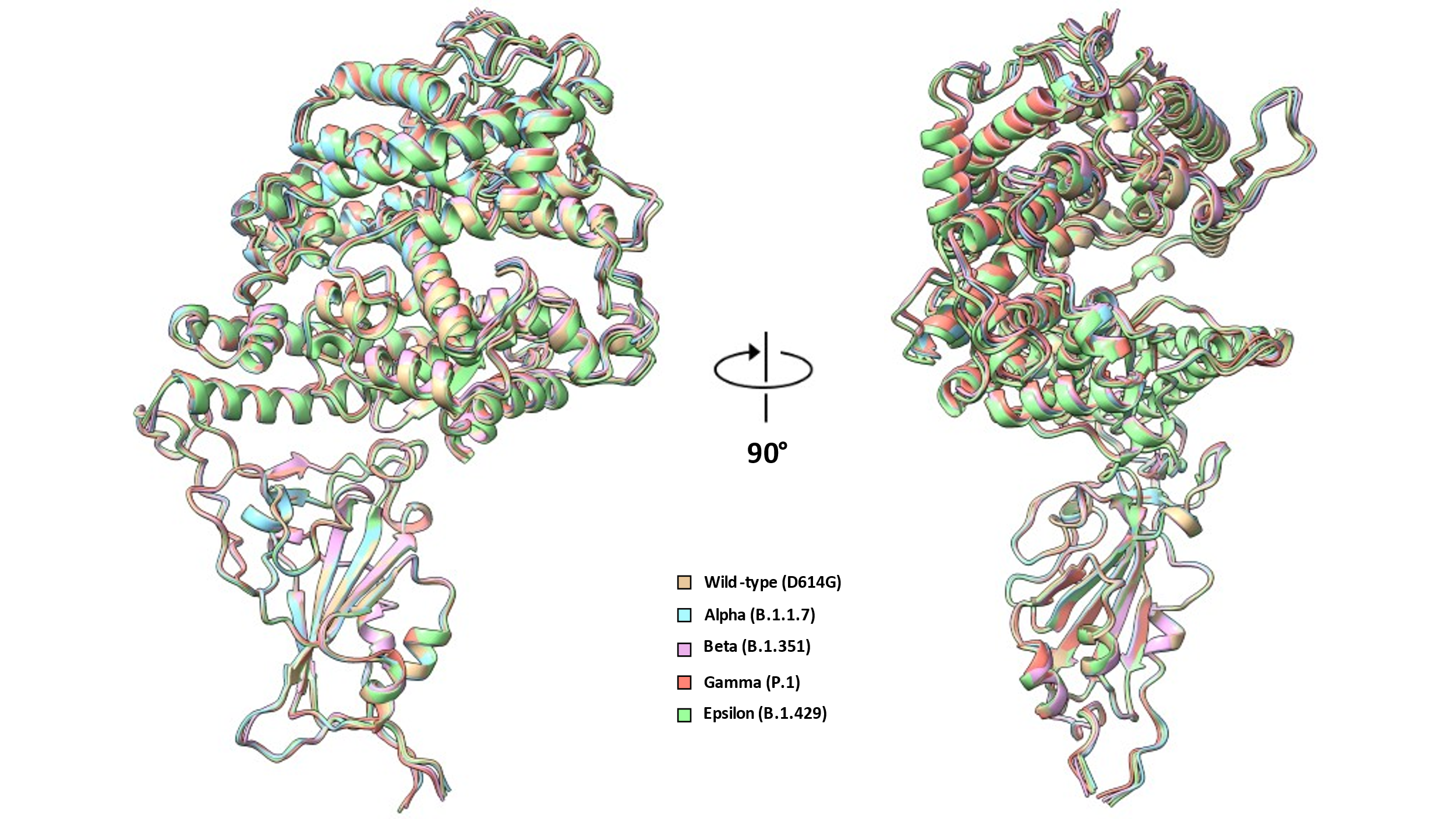

**Figure S22. Superposition of RBD-ACE2 local models.** Models were aligned using the RBD for the superposition.

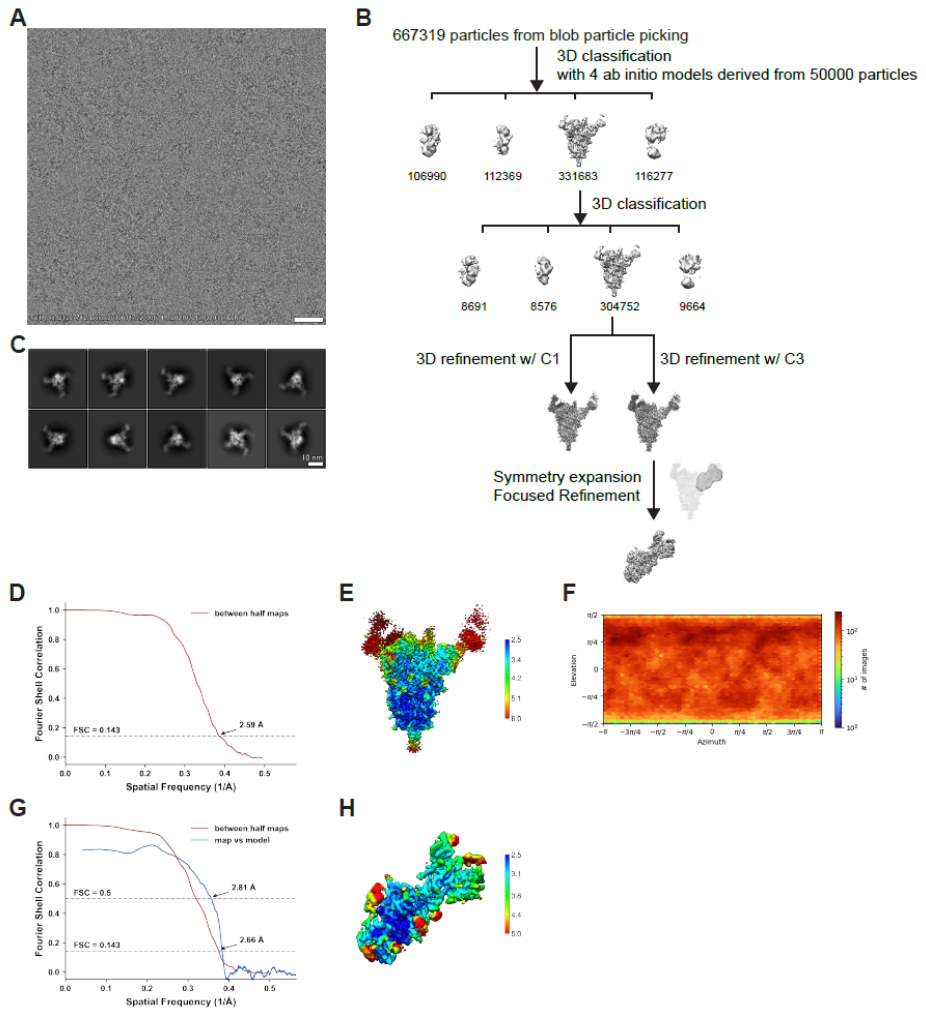

**Figure S23. Cryo-EM data processing and validation for complex of Gamma spike protein ectodomain and 4-A8 Fab.** **(A)** Representative cryo-EM micrograph. **(B)** Workflow of cryo-EM image processing. **(C)** Representative 2D classes. **(D-F)** FSC curves **(D)**, local resolution **(E)** and viewing direction distribution plot **(F)** of global refinement. **(G-H)** FSC curves **(G)** and local resolution **(H)** of focused refinement.

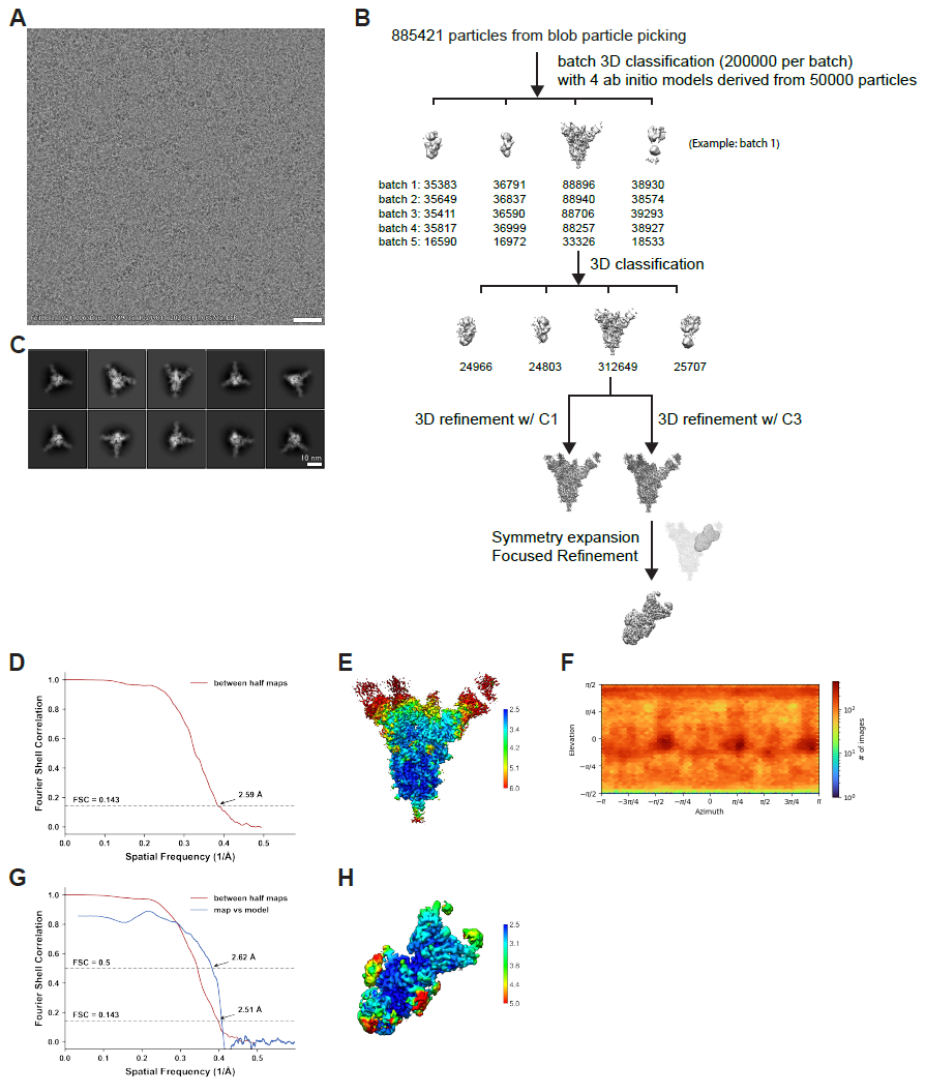

**Figure S24. Cryo-EM data processing and validation for complex of Gamma spike protein ectodomain and 4-8 Fab.** **(A)** Representative cryo-EM micrograph. **(B)** Workflow of cryo-EM image processing. **(C)** Representative 2D classes. **(D-F)** FSC curves **(D)**, local resolution **(E)** and viewing direction distribution plot **(F)** of global refinement. **(G-H)** FSC curves **(G)** and local resolution **(H)** of focused refinement.

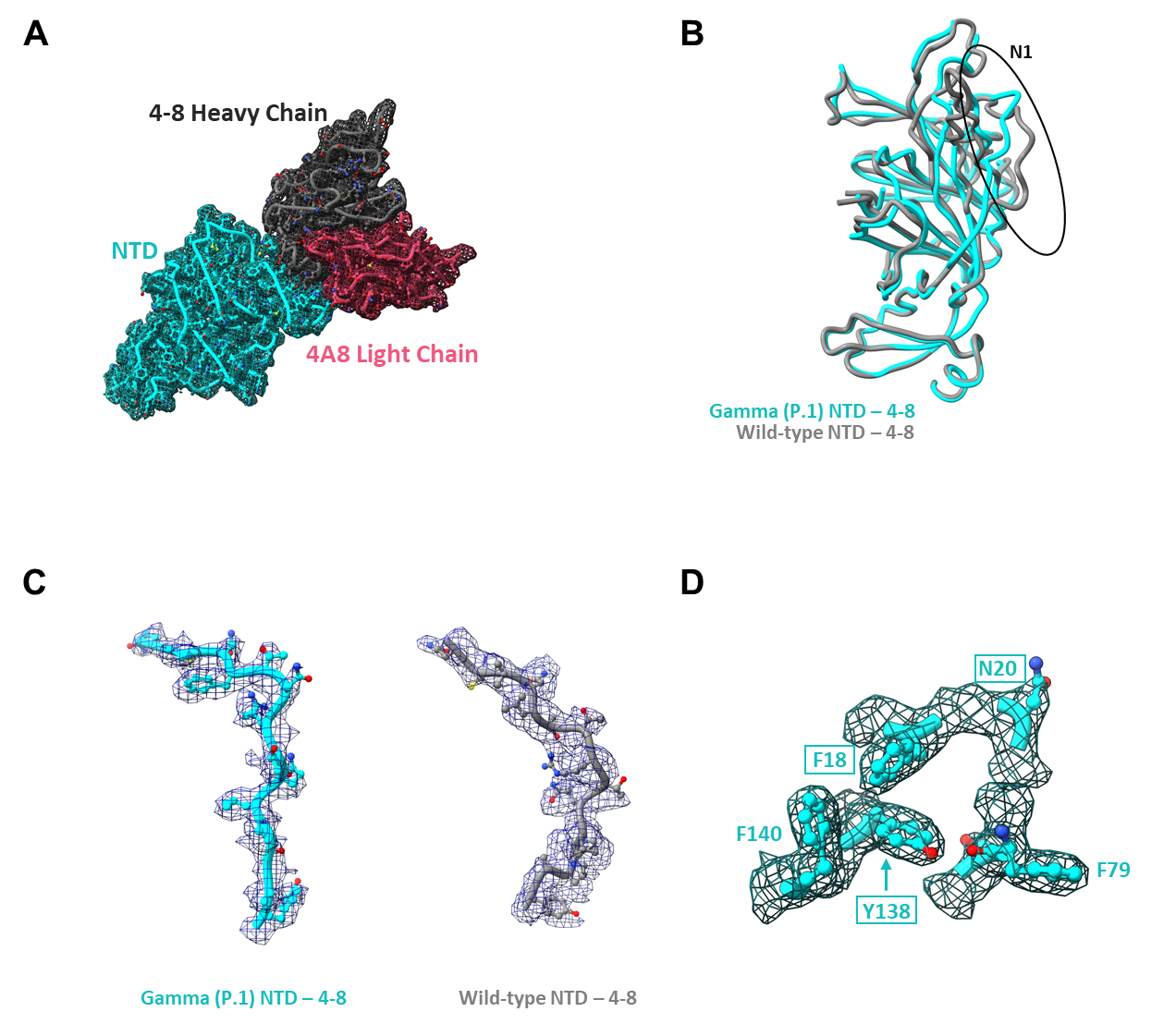

**Figure S25. Structure of the Gamma variant NTD bound by 4-8 reveals rearrangement of the N1 loop.** **(A)** Local cryoEM density map and model of the Gamma variant S protein bound to 4-8 at 2.51 Å **(B)** Superposition of 4-8-bound Gamma and wild-type NTD models showing N1 loop rearrangement. **(C)** Density and models for the N1 loops compared in panel (B). **(D)** Positioning of the L18->F, D138->Y, and T20->N mutations and adjacent residues in the Gamma NTD. **(D)** Superposition of residues shown in (C) with WT residues demonstrates steric incompatibilities. Areas of steric clashes are indicated by dashed ovals. Mutated residues are indicated as boxed labels. The wild-type – 4-8 model (PDB: 7LQV) was used for superpositions and is shown in grey throughout the figure.

**Table S1: CryoEM data collection, processing, refinement, and validation parameters for the structures reported in this publication.**

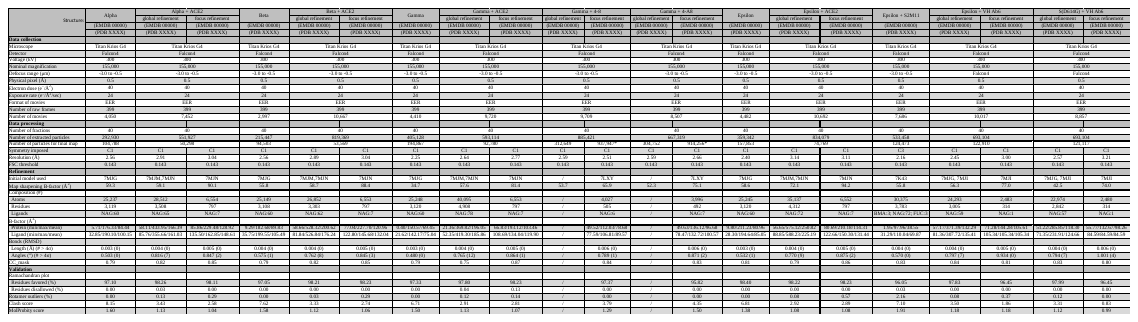
